# Rapid clonal selection within early hematopoietic cell compartments presages outcome to ivosidenib combination therapy

**DOI:** 10.1101/2024.12.30.630700

**Authors:** Sven Turkalj, Bilyana Stoilova, Angus J. Groom, Felix A. Radtke, Rabea Mecklenbrauck, Niels Asger Jakobsen, Curtis A. Lachowiez, Marlen Metzner, Batchimeg Usukhbayar, Mirian Angulo Salazar, Zhihong Zeng, Sanam Loghavi, Jennifer Marvin-Peek, Verena Körber, Farhad Ravandi, Ghayas Issa, Tapan Kadia, Vasiliki Symeonidou, Anne P. de Groot, Hagop Kantarjian, Koichi Takahashi, Marina Konopleva, Courtney D. DiNardo, Paresh Vyas

**Affiliations:** MRC Molecular Haematology Unit, Radcliffe Department of Medicine, Weatherall Institute of Medicine, University of Oxford, Oxford, United Kingdom; The University of Texas MD Anderson Cancer Center, Department of Leukemia, Houston, Texas, USA; The University of Texas MD Anderson Cancer Center, Department of Hematopathology, Houston, Texas, USA; Department of Haematology, Oxford University Hospitals NHS Trust, Oxford, United Kingdom

## Abstract

Acquired resistance to targeted non-intensive therapies is common in myeloid malignancies. Yet, key questions remain as to how rapidly resistant clones are selected by treatment and in which hematopoietic cell compartments clonal selection occurs. To address this gap, we studied clonal responses to ivosidenib + venetoclax ± azacitidine combination therapy in 8 patients with *IDH1*-mutant myeloid malignancy. Whilst all 8 patients initially responded to treatment, 6 relapsed and 2 remained in sustained remission for > 4 years. To study longitudinal clonal dynamics through hematopoietic differentiation, we performed high-sensitivity single-cell genotyping in index-sorted sequential patient samples. In all patients who relapsed, therapy-resistant clones were selected rapidly, within 1-3 treatment cycles, at times when hematopoiesis was still largely sustained by either normal or pre-leukemic cells. Selection of therapy-resistant clones preceded overt treatment failure by months or even years. Relapse was associated either with clones harboring newly-detected myeloid driver mutations or expansion of minor pre-existing clones that had reduced fitness prior to treatment. In both cases, resistant clones were selected within immature cell populations previously shown to contain leukemic stem cell (LSC) potential, preceding malignant expansion of these compartments by immunophenotyping. In contrast, in both patients remaining in remission, leukemic clones were eradicated and rapidly replaced by clonal and wild-type hematopoiesis. These observations suggest that, in patients treated with non-intensive ivosidenib combination therapy, rapid clonal selection occurs in populations with LSC potential, where failure to eliminate either genetically evolved or persistent leukemic clones ultimately leads to relapse.

**Key points:** – Rapid selection of leukemic clones occurs within small populations with LSC potential, months or years prior to relapse.

– Rapid eradication of leukemic clones leads to sustained remission in the context of ivosidenib combination therapy.

## Introduction

Acute myeloid leukemia (AML) is an aggressive malignancy resulting from differentiation delay/arrest and expansion of immature myeloid cells. Two intertwined hierarchies co-exist in AML^1^: a genetic clonal hierarchy, consisting of normal, pre-leukemic, and leukemic clones; and a hematopoietic hierarchy, where cells are arrayed through a continuum of hematopoietic differentiation.^2–7^ Whilst pre-leukemic mutations do not usually lead to differentiation arrest,^8–10^ subsequent genetic and/or epigenetic lesions impair differentiation of hematopoietic progenitors and/or precursors, enhance their self-renewal capacity, and impart functional leukemic stem cell (LSC) properties.^11,12^ Importantly, different leukemic driver mutations preferentially accumulate at distinct differentiation stages,^5,7,10,12–14^ which impacts therapy response.^13,15,16^

Most AML patients, including the ∼8-10% of patients presenting with recurrent somatic mutations in the isocitrate dehydrogenase 1 (*IDH1*) gene,^17–19^ are ineligible for curative intensive therapies.^20^ For this patient group, treatment options include: venetoclax (VEN) with azacitidine (AZA; VEN+AZA); low dose cytosine arabinoside^21–23^; and, for *IDH1*-mutated AML, the combination of the mutant IDH1 inhibitor ivosidenib (IVO) with AZA (IVO+AZA).^24^ Non-intensive treatment often produces an initial clinical response, but relapse occurs in most patients. Distinct and shared resistance mechanisms to VEN+AZA and IVO have been proposed.^25^ Shared mechanisms include mutations in genes encoding signaling molecules (e.g. *FLT3* and *RAS), TP53* mutations, and complex karyotype.^26–28^ Distinct resistance mechanisms include mutations in genes regulating apoptosis (e.g. *BAX*) in VEN-treated patients^29^ and *IDH2* mutations in IVO-treated patients (i.e. isoform switching).^27^ VEN+AZA resistance has also been attributed to reduced mitochondrial priming^30^ and higher anti-apoptotic BCL-xL and MCL-1 protein expression.^31^ FLT3 and RAS^28,32,33^ signaling, as well as monocytic differentiation,^34^ have been suggested to confer MCL-1-mediated VEN resistance.

To evaluate the efficacy of combining targeted therapy for mutant IDH1 and VEN-containing regimens, we conducted a phase Ib/II study of IVO+VEN±AZA in patients with *IDH1*-mutated myeloid malignancies^35^ (NCT03471260). The initial overall response rate (ORR) was 94% and the median follow-up was 24 months. In 6/9 patients with available samples who relapsed after achieving an initial response, relapse was associated with genetic evolution.^35^ Selected mutations occurred in genes encoding transcription factors and signaling molecules. However, the precise clonal and cellular context (i.e. stages of differentiation) where therapy-induced selection was occurring remained unclear, representing an important question for two reasons. First, both the genetic milieu of selected clones and the stage of differentiation arrest are associated with distinct clinical outcomes to therapy and *in vitro* drug screening.^13,15,16^ Second, myeloid driver mutations exert different molecular consequences at distinct differentiation stages and within distinct clonal contexts.^28,36–39^ To enable future mechanistic investigation of clone-specific vulnerabilities, the differentiation stage(s) where clonal selection occurs must be identified.

Furthermore, though the duration of response (DOR) varied substantially between patients (months to years), it remained unclear how quickly clonal selection leading to either resistance or sustained remission occurred and whether a correlation exists between the timing of selection and DOR. Finally, the longitudinal clonal architecture of patients with sustained response was not identified.

To address these questions, we selected 8 patients treated with IVO+VEN±AZA where an initial response was achieved and where sequential bone marrow (BM) samples were available. With longer follow-up, 6 patients relapsed while 2 achieved sustained remission for 48 and 57 months. In all sequential samples, we performed high-fidelity single-cell genotyping coupled with immunophenotypic and RNA-seq analysis,^38,40^ enabling the investigation of longitudinal clonal dynamics within hematopoietic differentiation.

## Methods

### Patients and sample processing

Patient samples were obtained at MD Anderson Cancer Center (NCT03471260 trial). Control BM samples were collected from age-/sex-matched individuals, with no hematologic malignancy (Nuffield Orthopaedic Centre, Oxford^38^). Informed consent was secured; studies were approved by the local ethics committees. Sample collection and processing was performed as described before.^35,38^

### Outcome parameters

Remission was defined according to 2022 ELN.^20^ Overall survival (OS) was defined from the first day of therapy to death or last follow-up. DOR was defined from the first documented therapy response (blast reduction/PR/CRi/CR) to treatment failure. Treatment failure was defined as relapse, disease progression, or the need to switch therapy.

### Bulk DNA sequencing

Detailed procedures are in Supplemental Methods. An 81-gene targeted Next Generation Sequencing (NGS) panel (Supplemental Table 1) was performed^35^; a 97-gene targeted NGS panel^38^ (Supplemental Table 1) was additionally performed for pts 4/9. Whole-exome sequencing (WES) was performed on pts 4/5/10/11 at baseline and pts 11/14 at relapse, using CD45^dim^ blasts and CD3^+^ T control cells.

### Cytogenetic analysis and molecular MRD assessment

Conventional karyotyping was performed.^35^ *IDH1* mutation was assessed by Sanger sequencing (10–20% sensitivity) and by NGS (1–2% sensitivity). Measurable residual disease (MRD) was assessed by flow cytometry (0.1-0.01% sensitivity).

### Flow cytometry and FACS

Primary sample FACS staining was performed^38^ (Supplemental Methods). Samples were stained with antibodies against CD38/CD10/CD117/CD45RA/CD123/CD90/CD34, an antibody lineage cocktail against CD2/CD3/CD4/CD8a/CD19/CD20/CD235, and 7-AAD. FACS analysis and single-cell FACS-sorting for TARGET-seq+ were performed. Single-cell index data was collected. Cells from the NOC156 control sample were sorted onto ∼10% of each plate. Flow cytometry data analysis on non-pre-enriched samples was performed using FlowJo v10.8.1 and R (v4.3.1).

### TARGET-seq+

Detailed procedures^38^ are in Supplemental Methods. Genotyping primers (Supplemental Table 3), designed with Primer BLAST,^41^ were tested as described.^38^ Transcriptome libraries were sequenced at 1 million reads/cell (except for pt5, sequenced at 40,000 reads/cell). Genotyping libraries were sequenced at 2,000 reads/amplicon/cell.

### TARGET-seq+ genotyping analysis

Detailed procedures are in Supplemental Methods. Briefly, the number of mutant and WT reads at each locus was computed using TARGET-seq,^40^ getITD,^42^ and custom pipelines. After filtering for cells with sufficient coverage, single-cell mutation calling was performed using single-cell variant allele frequency (scVAF) and mutant read number thresholds established by genotyping WT control BM cells.^1,37,38^

Where applicable, copy number alterations (CNAs) were inferred at single-cell level from RNA expression. Transcriptome reads were pre-processed with a custom pipeline,^38^ followed by quality filtering and normalization using Seurat (v5.0.1)^43^ and SingleCellExperiment (v1.24.0).^44^ inferCNV(v1.18.1),^45^ followed by a custom CNA calling strategy, was used to infer CNAs in single cells.

Single-cell mutation and CNA calls were merged. infSCITE^46^ was used to determine the statistically most likely phylogenetic tree. Using the infSCITE output, co-occurrence of somatic events was used to assign each cell to a clone, after correction for allelic drop-out. Each cell was assigned to an immunophenotypic population using surface markers. Clonal and immunophenotypic data were merged.

For pts 4/5/9/10/11/14/18, the size of each clone within all live, lineage-negative (Lin^-^) cells at each time point was inferred, considering the FACS size of each population, the number of cells sampled per population, and the clone sizes within each population.

### Quantification and statistical analysis

Statistical data analysis was performed using R (v4.3.1). Plots were generated using ggplot2 (v3.3.6) or FlowJo (v10.8.1). Statistical tests used and summary statistics are in each figure legend.

### Data availability

Bulk targeted NGS/WES/single-cell genotyping data are available at SRA (https://www.ncbi.nlm.nih.gov/bioproject/PRJNA1139762); accession number: PRJNA1139762. Single-cell RNA-seq data are available at GEO (https://www.ncbi.nlm.nih.gov/geo/query/acc.cgi?acc=GSE273135); accession number: GSE273135.

## Results

### Patient cohort and experimental strategy

8 patients (pts) treated with IVO+VEN±AZA were studied (Figure 1A; Table 1). Pts 11/5/20/14 were newly diagnosed, whereas pts 9/4/18/10 had relapsed disease. Treatment used was: VEN on days 1–14; IVO 500 mg continuously, beginning on cycle (C) 1 day 15; and, in pts 18/20, AZA 75mg/m^2^ for 7 days was also administered. pt20 received IVO+VEN±AZA for two cycles, followed by IVO monotherapy, after experiencing prolonged cytopenia in C1-2. After a median follow-up of 40.3 months (data cut-off 2023-12-14), median DOR was 18.5 months, median time on treatment was 20.3 months, and median OS was 45.5 months. At the time of data cut-off, pts 10/18 remained on treatment.

**Figure 1.**
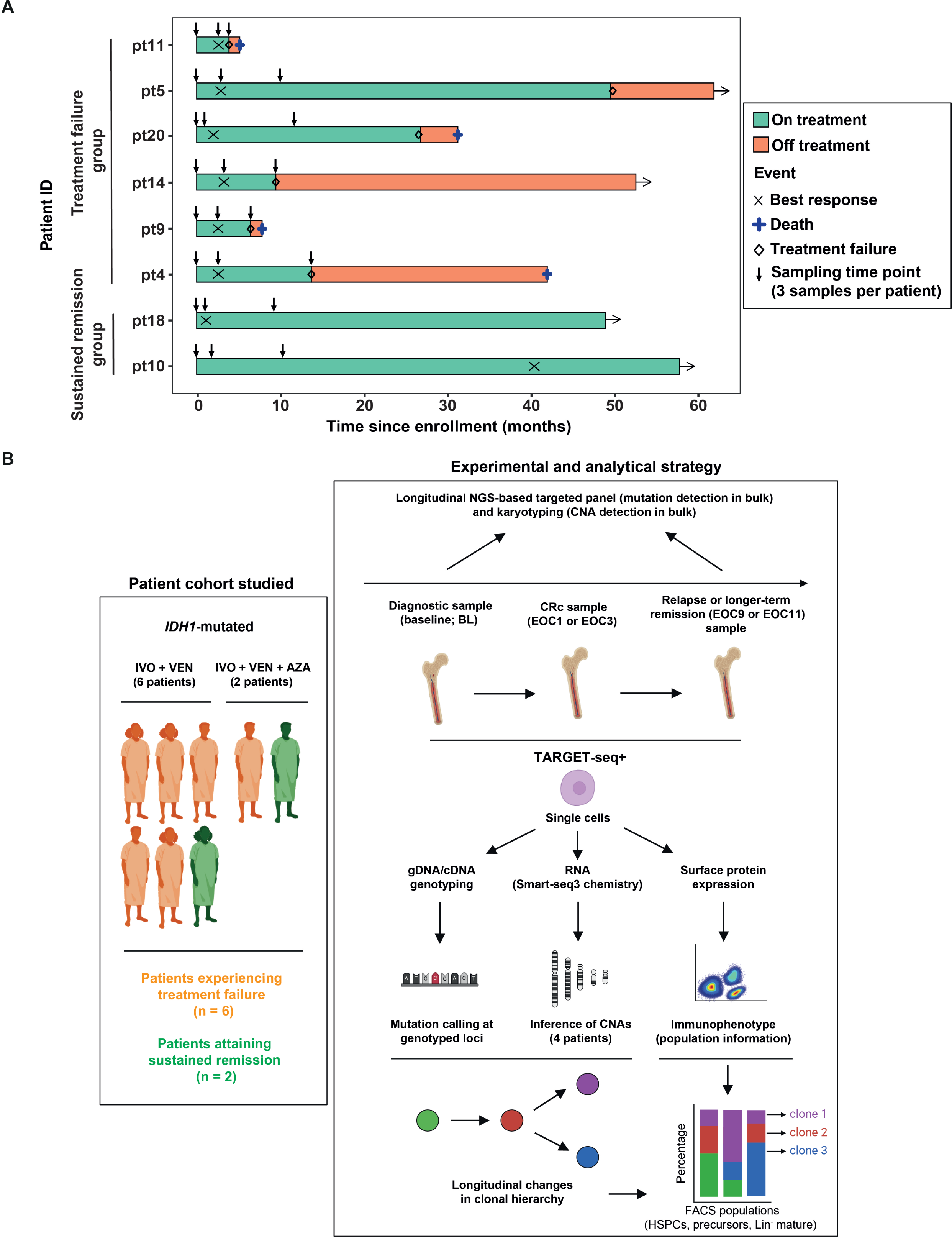
Patient cohort and experimental strategy. (A) Swimmer plot showing response to treatment. “X” indicates time point of first best treatment response. Treatment failure (black rhombus) is defined as either disease progression or treatment failure. Vertical arrows indicate sampling time points. Patients are separated into two groups according to treatment response. Patients are shown in the order in which they are mentioned in the text. (B) Left, 6 patients were treated with IVO and VEN (pts 4, 5, 9, 10, 11, and 14); 2 patients with IVO, VEN, and AZA (pts 18 and 20). 6 patients progressed (orange); 2 patients remain in remission (green). Sequential samples were studied from three time points: (a) pre-treatment (baseline, BL); (b) early treatment response (end of cycle (EOC) 1 for pts 18 and 20, EOC3 for pts 4, 5, 9, 10, 11, and 14); (c) relapse (pts 4, 9, 11, and 14), EOC9 (pt18), or EOC11 (pts 5, 10, and 20). Samples were studied by TARGET-seq+ to obtain single-cell genotype (somatic mutations obtained through targeted genotyping; copy number alterations inferred through scRNA-seq) and immunophenotype. Clonal and immunophenotypic information was then merged for each single cell.

**Table 1.**

| Characteristic | pt11 | pt5 | pt20 | pt14 | pt9 | pt4 | pt18 | pt10 |
| --- | --- | --- | --- | --- | --- | --- | --- | --- |
| Age at time of enrollment (years) | 65 | 84 | 63 | 80 | 77 | 68 | 75 | 72 |
| Sex | Female | Female | Male | Female | Male | Male | Male | Female |
| Diagnosis | AML | AML | AML | AML* | R/R AML | MDS/MPN | sAML | R/R AML |
| ECOG | 1 | 1 | 1 | 2 | 2 | 1 | 0 | 1 |
| Previous therapies | N/A | N/A | N/A | N/A | DEC | AZA; FT2102 | AZA | AZA; IA/LY251092;<br>DEC; AZA/FT-2102;<br>alloHCT |
| Previous IDH1<sup>MUT</sup> inhibitor | No | No | No | No | No | Yes | No | Yes |
| WBC at baseline (10<sup>3</sup>/ml) | 11 | 1.5 | 1.2 | 3.7 | 0.6 | 4.6 | 1.8 | 1.9 |
| Hb at baseline (g/dl) | 10.1 | 9.4 | 10.1 | 8 | 8.9 | 11.2 | 8.7 | 7.7 |
| Platelets at baseline (10<sup>3</sup>/ml) | 53 | 185 | 6 | 57 | 5 | 160 | 74 | 12 |
| Blast Count at baseline (%) | 28 | 47 | 79 | 16 | 61 | 12 | 10 | 23 |
| IDH1 mutation at baseline (VAF [%]) | R132H (29) | R132H (22) | R132C (42) | R132H (32) | R132C (31) | R132H (38) | R132H (17) | R132S (18) |
| Treatment | IVO 500mg +<br>VEN 800mg | IVO 500mg +<br>VEN 400mg | IVO 500mg +<br>VEN 400mg +<br>AZA 75mg/m<sup>2</sup> | IVO 500mg +<br>VEN 800mg | IVO 500mg +<br>VEN 800mg | IVO 500mg +<br>VEN 400mg | IVO 500mg +<br>VEN 400mg +<br>AZA 75mg/m<sup>2</sup> | IVO 500mg +<br>VEN 800mg |
| Time on treatment (months) | 3.9 | 49.6 | 26.8 | 9.5 | 6.5 | 13.7 | 49.0 | 57.8 |
| Treatment failure | Yes | Yes | Yes | Yes | Yes | Yes | No | No |
| Duration of response (months) | 3.4 | 47.5 | 24.3 | 6.7 | 3.9 | 12.7 | 48.1 | 57.0 |
| Best response | CR | CR | MLFS | CR | CR | CR | CRi | CR |
| Overall survival (months) | 5.1 | 62.0 | 31.3 | 52.6 | 7.8 | 42 | 49.0 | 57.8 |
| Survival status | Deceased | Alive | Deceased | Alive | Deceased | Deceased | Alive | Alive |
| Follow-up treatment | FIA/gilteritinib | LEN | VEN/HHT | alternating CLAD/LDAC<br>and DEC/VEN | gilteritinib/VEN | CLAD/LDAC | N/A | N/A |

AML: De novo acute myeloid leukemia; sAML: Secondary AML; MDS/MPN: Myelodysplastic/myeloproliferative neoplasm; R/R AML: Relapsed/refractory AML; ECOG: Eastern Cooperative Oncology Group Performance Status Scale; Hb: Hemoglobin; WBC: White blood cell count; VAF: Variant allele frequency; IVO: Ivosidenib; VEN: Venetoclax; AZA: Azacitidine; CR: Complete remission; CRi: CR with incomplete hematologic recovery; MLFS: Morphologic leukemia-free state; DEC: Decitabine; LEN: Lenalidomide; CLAD: Cladribine; LDAC: Low-dose cytarabine; IA: Idarubicin/cytarabine; alloHCT: Allogeneic stem cell transplantation; FIA: Fludarabine/idarubicin/cytarabine; HHT: Homoharringtonine.
\* pt14 has originally been classified as MDS/MPN (Lachowiez et al., 2023). According to ICC 2022 and the 5<sup>th</sup> edition of the WHO classification, for this study, pt14 was re-classified as AML with mutated *NPM1*, due to the presence of high *NPM1* VAF and blasts > 10% at baseline.

Our approach is illustrated in Figure 1B. All samples were karyotyped (Supplemental Table 1A) and screened with an 81-gene NGS panel (Supplemental Table 1B) at baseline and relapse (Supplemental Tables 1D, 1F).^35^ We additionally performed a 97-gene targeted NGS panel at 822x (for pts 4/9; Supplemental Table 1C) and WES at 100x (for pts 4/5/10/11 at baseline and pts 11/14 at relapse; Supplemental Tables 1E and 1G). MRD was assessed by flow cytometry (threshold 0.1-0.01%; Supplemental Table 1I).

We first established the sizes of immunophenotypic normal BM mononuclear cell (BMMNC) lineage-negative (Lin^-^) compartments (all populations except for mature lymphoid and erythroid cells) in 10 normal human BM samples (Supplemental Figure 1A; Supplemental Table 2A). We defined the sizes of the following BM compartments: Lin^-^CD34^+^ hematopoietic stem and progenitor cells (HSPC), Lin^-^CD34^-^CD117^+^ granulocyte-macrophage (GM) precursors, and Lin^-^CD34^-^CD117^-^ mature myeloid cells. We then analyzed these compartments in patient samples (Supplemental Figures 1B–1D; Supplemental Table 2B).

Next, we index-sorted single cells from the HSPC, GM precursor, and mature myeloid compartments, from: (a) baseline; (b) early response; and (c) either treatment failure or long-term remission time points. Single-cell genotype, immunophenotype, and transcriptome were obtained. Clonal identities of single cells were assigned using patient-specific mutational panels (Supplemental Tables 3A–3D) and CNA inference from RNA-seq (Supplemental Tables 3E-3G). We did not interrogate specific *CEBPA* (pts 11/9) and *RUNX1* (pt9) mutations, as we could not design suitable genotyping primers. For pts 4/5/9/11/14, we did not genotype mutations displaying low VAFs (≤ 5%) at both baseline and relapse, hypothesizing they are unlikely to substantially contribute to relapse. A total of 14,383 single cells were assigned to clones. Finally, we integrated clonal status and immunophenotypic profile for each cell.

### Rapid selection of newly-detected resistant clones occurs within immunophenotypic leukemic stem cells, months or years prior to treatment failure

In 6 patients who relapsed, the time from initial response to relapse varied from 3 months (pt11) to ∼4 years (pt5). We identified 2 different patterns of clonal selection. Pts 11/5/20 relapsed with clones undetected prior to treatment, with relapse occurring months (pt11) or years (pts 5/20) after treatment initiation. Conversely, in pts 14/9/4, minor clones detected at trial entry expanded upon therapy exposure and became dominant at relapse. We first report on patients with newly-detected clones.

Pt11 had a DOR of 3.4 months. Relapse was associated with a genetically evolved clone, not detected at baseline, which was already selected by end of cycle 3 (EOC3). Pre-treatment, *DNMT3A* (D), *NPM1* (N), *IDH1* (I), and *NRAS* (Nr) mutations were detected (Figures 2A-2B; Supplemental Figures 2A–2B show single-cell genotyping thresholds). Additionally, a *CEBPA* mutation was detected by bulk NGS but was not genotyped at single-cell level (outlined above). Single-cell analysis revealed two dominant clones at baseline, DNI and DNINr (Figure 2B; Supplemental Table 4A). Both clones dominated the GM precursor compartment (the most expanded population) but were also detected within lymphoid-primed multipotent progenitors (LMPP) and granulocyte-monocyte progenitors (GMP; Figure 2C). All three compartments contain LSC potential.^11,12^ By EOC3, the patient had achieved CR, with the BM largely composed of wild type (WT) mature myeloid cells; however, we also detected a genetically evolved clone harboring an additional *FLT3*-ITD (F) mutation (DNINrF clone). Already by EOC3, the DNINrF clone comprised ∼90% of the small-sized LMPP, GMP, and GM precursor BM compartments (Figures 2A–2C). Subsequently, the DNINrF clone led to rapid relapse post-C3.

**Figure 2.**
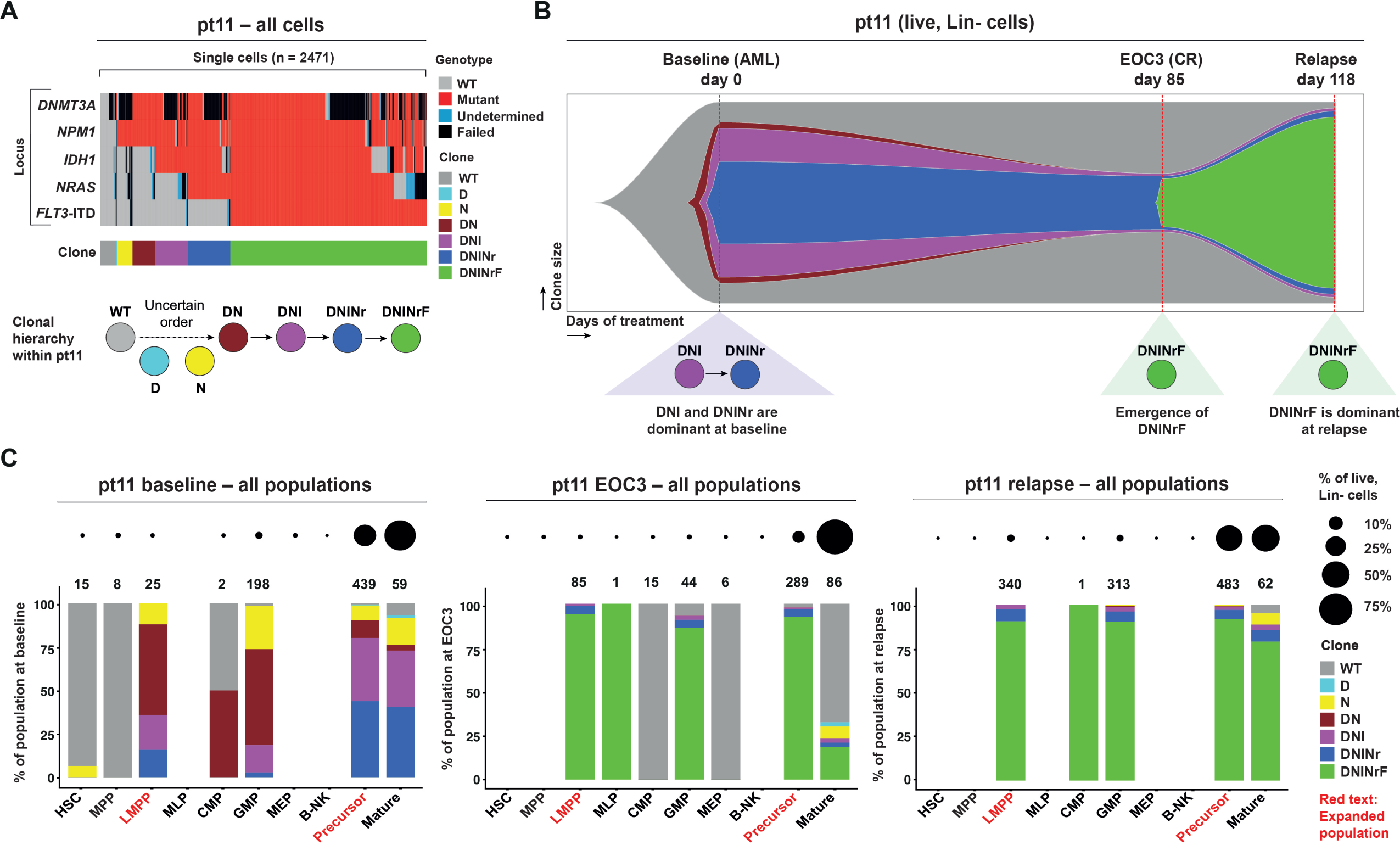
Selection of a *FLT3-ITD* resistant clone occurs within 3 months of treatment within immunophenotypic leukemia stem cells. (A) Top, raster plot of single-cell genotyping for pt11, within all cells assigned to a clone (n = 2471 cells). Each row represents a genotyped locus (left). The genotyped mutations are: *DNMT3A*-N757K, *NPM1*-W288fs*12, *IDH1*-R132H, *NRAS*-G12C, *FLT3*-ITD. Each column is a single cell. Colors indicate whether a cell was called wild type (WT, gray), mutant (red), undetermined (blue), or whether genotyping failed (Failed, black). Middle, clonal assignment for each cell. Color legend on the right. Due to suboptimal genotyping at the *DNMT3A* locus, the order of acquisition of the *DNMT3A* and *NPM1* mutations was uncertain. Clone abbreviations are: WT: wild type (gray). D: *DNMT3A* mutant (cyan). N: *NPM1* mutant (yellow). DN: *DNMT3A* and *NPM1* mutant (brown). DNI: *DNMT3A*, *NPM1*, and *IDH1* mutant (purple). DNINr: *DNMT3A*, *NPM1*, *IDH1*, and *NRAS* mutant (blue). DNINrF: *DNMT3A*, *NPM1*, *IDH1, NRAS*, and *FLT3* mutant (green). Bottom, most likely clonal structure for pt11 (Methods). (B) Longitudinal changes in clone size (y-axis) within all live, Lineage-negative (Lin^-^) cells (consisting of HSPC, GM precursor, and mature myeloid cells) in pt11, showing the baseline, end of cycle 3 (EOC3), and relapse time points (x-axis). Clone color-coding as in (A). The x-axis is proportional to the time (in days) relative to baseline. CR: complete remission; AML: *de novo* AML. Clone size inference within live, Lin^-^ cells detailed in Methods (computed frequencies and confidence intervals in Supplemental Table 4). (C) Clonal composition (indicated as percentage, y-axis) of each cell population sorted by FACS (x-axis; population abbreviations as in Supplemental Figure 1A) in pt11 at baseline (left), EOC3 (middle), and relapse (right). Clone color-coding as in (A). Numbers of profiled cells per population are indicated above the bars. The size of the black dots is proportional to the percentage of a given population within all live Lin^-^ cells at a given time point, prior to any FACS enrichment (legend on the right). The red text marks populations exceeding the mean size of the respective healthy control BM populations by more than four standard deviations.

In 2 other patients (pts 5/20), newly-detected clones were also selected early but frank treatment failure ensued with much slower kinetics. Pre-treatment, pt5 had del(5)(q22q35) (−5q) and *RUNX1* (R), *ASXL1* (A), *IDH1* (I), and *TET2* (T) mutations (Figures 3A–3B; Supplemental Figures 3A–3D). The (−5q)RAI clone, dominant at baseline, was eradicated by EOC3 (Figures 3A–3B; Supplemental Table 4B), when the patient was in CR and mature myeloid cells were mainly WT. However, at EOC3, the ancestral (−5q) clone had persisted in ∼1% of all BM Lin^-^ cells. Moreover, a previously undetected clone harboring the *SF3B1*-K700E mutation (clone (−5q)S) was present in < 0.1% of all BM Lin^-^ cells (Figures 3A–3B). Interestingly, the (−5q)S clone was detected in 4/12 of immunophenotypic hematopoietic stem cells (HSCs; Figure 3C), suggesting this clone originated within HSCs. In parallel, an independent *TET2*-mutated clone (T clone), already detected at baseline, also persisted. By EOC11, when the patient was still in CR, all 3 clones ((−5q), (−5q)S, and T) had expanded. However, whereas the T clone differentiated efficiently into mature cells, (−5q) and (−5q)S clones were dominant in HSCs (70% of HSC) and megakaryocyte-erythroid progenitors (MEP; 56% of MEP; Figure 3C). Notably, AML subtypes with erythroid differentiation rely on BCL-xL for their survival.^47^ Pt5 was MRD-negative by flow cytometry both at EOC3 and EOC11 (Supplemental Table 1I). 40 months after EOC11, pt5 progressed to myelodysplasia (MDS) with the (−5q)S clone (Supplemental Tables 1A and 1F), leading to lenalidomide monotherapy being initiated.

**Figure 3.**
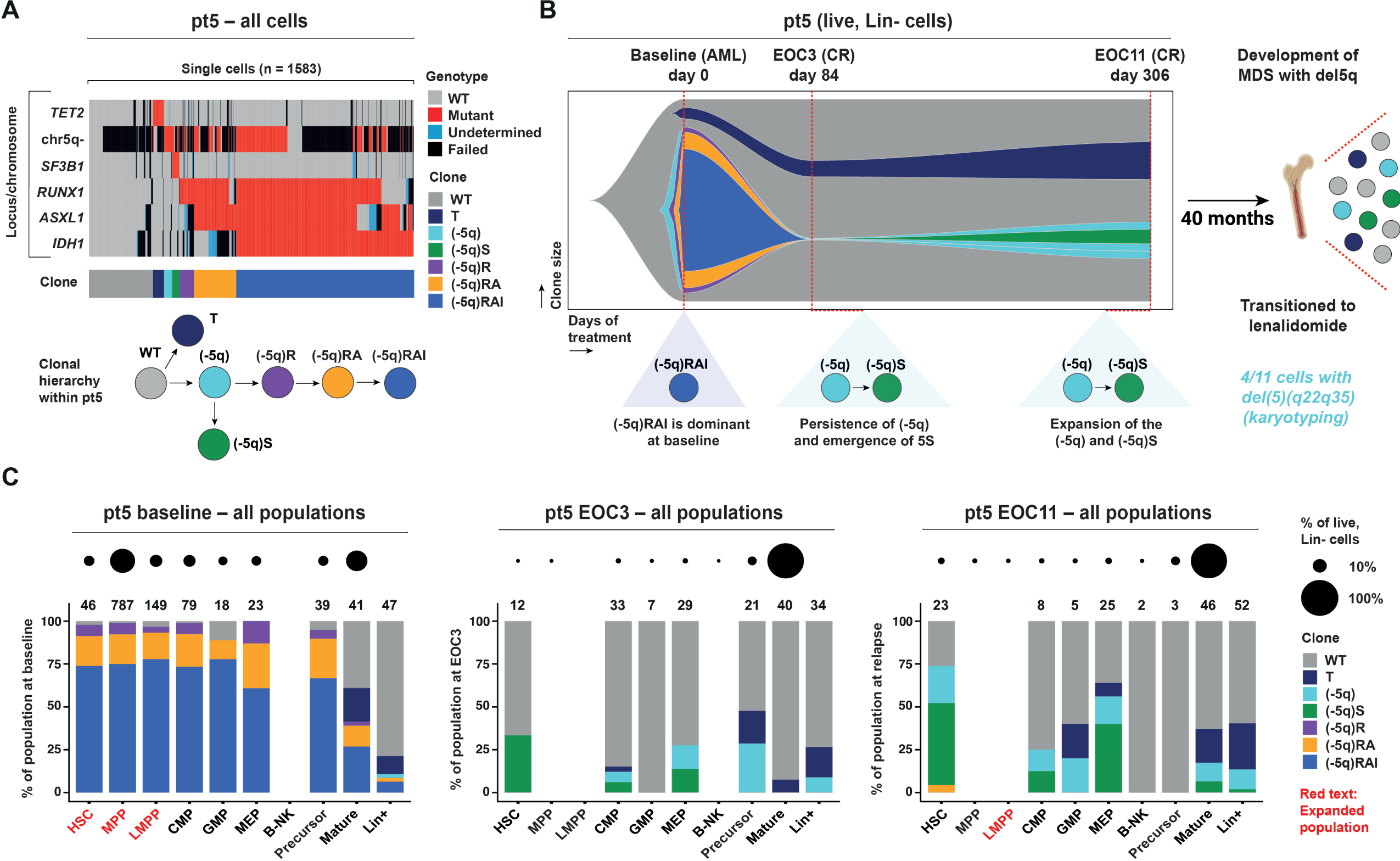
In pt5, selection of resistant clones within immunophenotypic stem cells occurred years prior to treatment failure. (A) Same panel structure as in Figure 2A but for pt5 (n = 1583 cells). The genotyped mutations are: *TET2*-K889*, *SF3B1*-K700E, *RUNX1*-R162K, *ASXL1*-H630fs, *IDH1*-R132C. inferCNV was used for single-cell CNA calling at chromosome 5 (del(5)(q22q35) was identified by karyotyping; referred to as chr5q-). Cells black for chr5q-did not pass scRNA-seq quality control (QC; Methods). Clone abbreviations are: WT: wild type (gray). T: *TET2* mutant (dark blue). (−5q): chr5q-(cyan). (−5q)S: chr5q- and *SF3B1* mutant (green). (−5q)R: chr5q- and *RUNX1* mutant (purple). (−5q)RA: chr5q-, *RUNX1*, and *ASXL1* mutant (orange). (−5q)RAI: chr5q-, *RUNX1*, *ASXL1,* and *IDH1* mutant (blue). (B) Same panel structure as in Figure 2B but for pt5, showing the baseline, EOC3, and EOC11 time points. Clone color-coding as in (A). Right, 40 months following EOC11, pt5 developed MDS associated with del(5)(q22q35) and *SF3B1*-K700E. (C) Same panel structure as in Figure 2C but for pt5, across baseline, EOC3, and EOC11 time points. Clone color-coding as in (A). In this patient, Lineage-positive (Lin^+^) cells were also genotyped.

Pt20 had a DOR of 24.3 months. Pre-treatment, two *TET2* (T), and *ASXL1* (A), *SRSF2* (S), *IDH1* (I), and *BRAF* (B) mutations were detected within a complex clonal structure (Supplemental Figures 4A–4C). The TTASI clone, dominant pre-treatment, was eradicated by EOC1 (BM blast reduction from 79% to 6%) and replaced by hematopoiesis from its parent clones (TTA/TTAB/TTAS; Supplemental Figures 4C–4D). In parallel, a newly-detected *RUNX1*-Y414fs*186 (R) mutation was selected within a minor terminal TTASR clone in the small-sized common myeloid progenitor (CMP) compartment, previously shown to have LSC potential.^48^ By EOC11, when the patient was in morphological leukemia-free state (MLFS), the TTASR/TTAS clones had expanded in the CMP, MEP, and GM precursor compartments, whereas TTAB/TTA/TT clones were more prominent in mature myeloid cells (Supplemental Figure 4D). 15 months later, the expansion of a primitive myeloid population gave rise to AML with the *RUNX1*-Y414fs* mutation (VAF of 69% in BMMNCs; Supplemental Table 1F),^35^ consistent with the hypothesis that defective differentiation of the TTASR clone led to treatment failure. Although this patient remained MRD-positive by flow cytometry, the dynamics of leukaemia-associated immunophenotypes (LAIP) did not reflect the kinetics of an expanding clone (Supplemental Table 1I).

**Figure 4.**
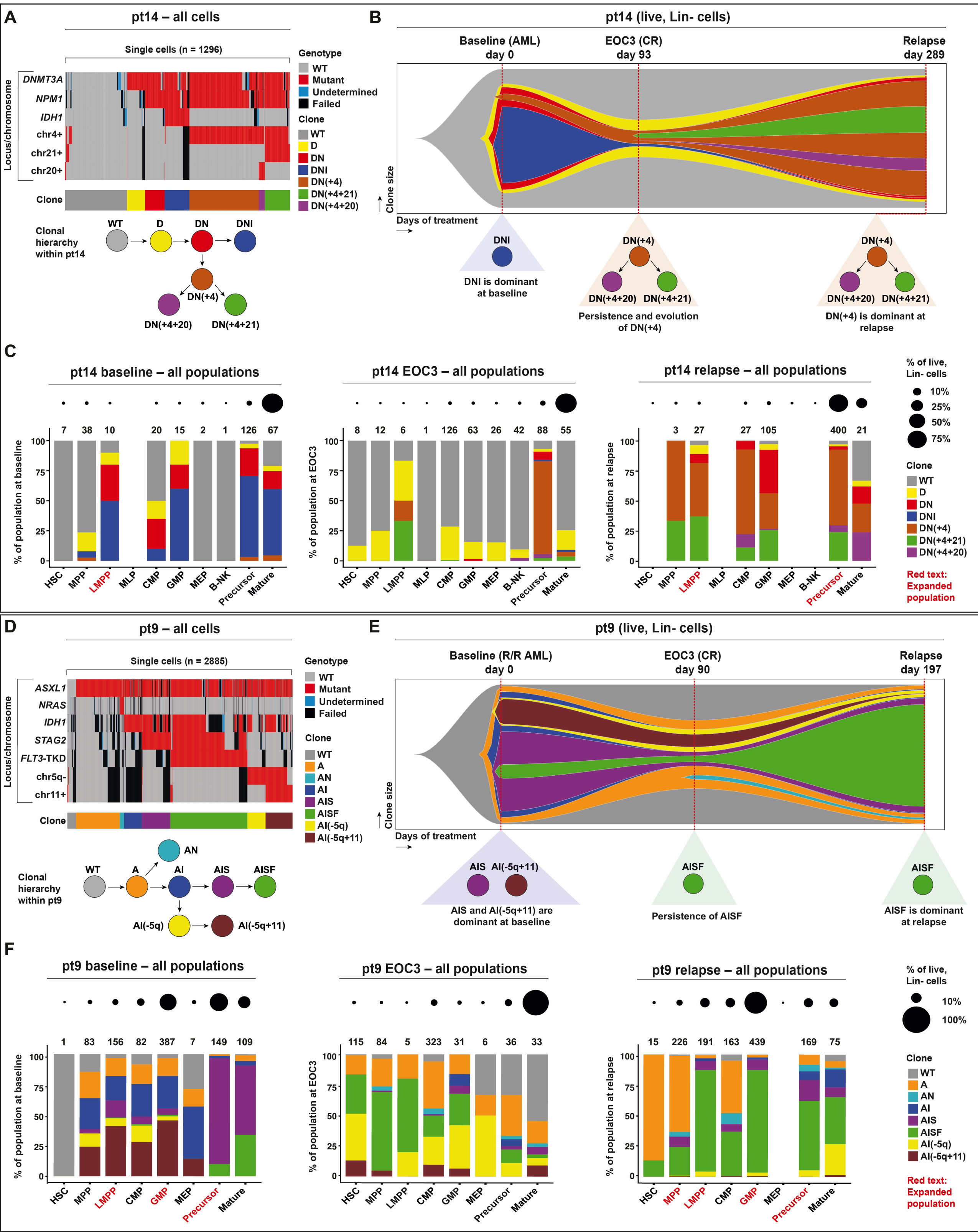
Minor pre-existing resistant clones expand within the first 3 treatment cycles within immunophenotypic LSCs. (A) Same panel structure as in Figure 2A, but for pt14 (n = 1296 cells). The genotyped mutations are: *DNMT3A*-G543C, *NPM1*-W288fs*12, *IDH1*-R132H. inferCNV was used for single-cell copy number alteration (CNA) calling at chromosomes 4, 21, and 20 (+4, +20, and +21 were identified by karyotyping - referred to as chr4+, chr20+, and chr21+, respectively). Clone abbreviations are: WT: wild type (gray). D: *DNMT3A* mutant (yellow). DN: *DNMT3A* and *NPM1* mutant (red). DNI: *DNMT3A*, *NPM1*, and *IDH1* mutant (blue). DN(+4): *DNMT3A*, and *NPM1* mutant with chr4+ (brown). DN(+4+21): *DNMT3A* and *NPM1* mutant with chr4+ and chr21+ (green). DN(+4+20): *DNMT3A* and *NPM1* mutant with chr4+ and chr20+ (purple). (B) Same panel structure as in Figure 2B but for pt14. Clone color-coding as in (A). (C) Same panel structure as in Figure 2C but for pt14. (D) Same panel structure as in (A) but for pt9 (n = 2885 cells). The genotyped mutations are: *ASXL1*-Q512*, *NRAS*-G13R, *IDH1*-R132C, *STAG2*-R1033*, *FLT3*-D835H. Additionally, inferCNV was used for single-cell copy number alteration (CNA) calling at chromosomes 5 and 11 (del(5)(q22q33) and +11 were identified by karyotyping; referred to as chr5q- and chr11+, respectively). Cells black for chr5q- and chr11+ did not pass scRNA-seq QC (Methods). Clone abbreviations are: WT: wild type (gray). A: *ASXL1* mutant (orange). AN: *ASXL1* and *NRAS* mutant (cyan). AI: *ASXL1* and *IDH1* mutant (blue). AIS: *ASXL1*, *IDH1*, and *STAG2* mutant (purple). AISF: *ASXL1*, *IDH1*, *STAG2,* and *FLT3* mutant (green). AI(−5q): *ASXL1* and *IDH1* mutant with chr5q-(yellow). AI(−5q+11): *ASXL1* and *IDH1* mutant with chr5q- and chr11+ (brown). (E) Same panel structure as in (B) but for pt9. Clone color-coding as in (D). R/R AML: relapsed/refractory AML. (F) Same panel structure as in (C) but for pt9. Clone color-coding as in (D).

Overall, in all 3 patients, therapy-resistant clones, undetected prior to therapy, were selected within 1–3 treatment cycles and expanded within small-sized LSC populations. Clonal selection within LSCs occurred months or years prior to treatment failure.

### Minor pre-existing clones expand under therapeutic pressure within immunophenotypic LSCs months prior to relapse

In 3 other patients (pts 14/9/4), therapy resistance arose from complex clonal adaptation to treatment.

In pt14, *DNMT3A* (D), *NPM1* (N), and *IDH1* (I) mutations, and trisomy 4 (+4), were detected at trial entry (Figures 4A–4B; Supplemental Figures 5A–5D). The DNI clone was dominant at baseline (Figure 4B; Supplemental Table 4C), whereas the DN(+4) clone was detected at similarly low levels in multipotent progenitors (MPP), GM precursors, and mature myeloid cells (Figure 4C). By EOC3 (CR), the DNI clone was nearly eradicated, and hematopoiesis was largely WT (Figures 4B–4C). However, the DN(+4) clone was now detectable in the LMPP compartment and dominated specifically the GM precursor compartment, despite both populations being small-sized (Figure 4C). Although the DN(+4) clone dominated GM precursors, its contribution to mature myeloid cells was very limited, suggesting it differentiated abnormally (Figure 4C). 6 months later, at relapse, the DN(+4) clone dominated the LMPP compartment and was still the major contributor to the expanded GM precursor compartment. Although the DN(+4) clone was detected in mature myeloid cells, its frequency was higher in HSPCs/GM precursors compared to more mature myeloid cells (Supplemental Figure 5E). Though two smaller clones evolved from DN(+4) with trisomy 20 or 21, the dominant clone at relapse was DN(+4).

In pt9, who had prior decitabine therapy, *ASXL1* (A), *IDH1* (I), *STAG2* (S), and *FLT3* tyrosine kinase domain (*FLT3*-TKD, F) mutations, del(5)(q22q33) (−5q), and trisomy 11 (11+), were detected at baseline (Figures 4D–4E; Supplemental Figures 6A–6E). Pre-treatment, AIS and AI(−5q+11) clones were dominant (Figure 4E; Supplemental Table 4D) in different BM compartments (Figure 4F). A smaller *FLT3*-TKD-mutated AISF clone was present within GM precursors and differentiated into mature myeloid cells (Figure 4F). By EOC3, when the patient was in CR, 73% of mature myeloid cells were either WT or harbored a single *ASXL1* mutation. However, the AISF clone had gained an advantage within multiple small-sized HSPC compartments, including populations with LSC potential (LMPP/GMP),^11^ while its contribution to mature myeloid cells was now minimal (Figure 4F). Relapse ensued months later, characterized by the expanded GMP (now the dominant population) and, to a lesser extent, LMPP compartments. While the AISF clone dominated GMP/LMPP compartments (Figure 4F), its frequency was lower in mature myeloid cells (Supplemental Figure 6F), suggesting impaired differentiation of this clone. Of note, 2 *CEBPA* mutations and 1 *RUNX1* mutation were detected by bulk NGS at relapse, at VAF > 10% (Supplemental Table 1F); we were technically unable to genotype these loci at single-cell level, limiting the clonal hierarchy dissection at relapse.

Similar but more subtle clonal dynamics were observed in pt4 (Supplemental Figure 7), who had progressed following treatment with AZA and the mutant IDH1 inhibitor olutasidenib. At baseline, the expanded CMP compartment was dominated by a clone with *SRSF2* (S), *IDH1* (I), *ASXL2* (A), and *JAK2* (J) mutations (SIAJ clone). Although the SIAJ clone persisted through treatment, it was reduced in size compared to baseline (Supplemental Figures 7D-7F). Conversely, two clones which evolved from the SIA clone by acquiring multiple CNAs (SIA(−20q-7) and SIA(−20q+1) clones), both minor at baseline, expanded by EOC3 (Supplemental Figure 7F). At relapse, which ensued ∼1 year later, both clones accumulated within the re-expanded CMP and other HSPC compartments. However, relapse was highly polyclonal, with expanded HSPCs compartments containing multiple clones that did not mature efficiently into myeloid cells (Supplemental Figures 7G–7H).

Altogether, in these 3 patients, minor pre-existing clones expanded rapidly upon therapy exposure within immunophenotypic LSCs. Despite their small size at remission, resistant clones accumulated within residual cells with LSC potential and differentiated inefficiently into mature myeloid cells.

### Sustained treatment response is associated with eradication of leukemic clones and selection of clonal hematopoiesis with a pre-leukemic driver

Finally, we report on 2 patients (pts 18/10) who remained in remission for 4.1 and 4.8 years, respectively, at the point of data cut-off.

Pt18 relapsed after AZA monotherapy prior to enrollment (Supplemental Figure 8A). The BM at baseline contained *RUNX1* (R), *IDH1* (I), *SRSF2* (S), and *ASXL1* (A) mutations (Figures 5A–5B; Supplemental Figures 8B–8C). Most of the BM was composed of the RIS/RI clones (Figure 5B; Supplemental Table 4F), dominant within expanded LMPP/GMP compartments (Figure 5C). A minor independent *ASXL1*-mutated clone was also detected. By EOC1, when pt18 achieved CR with incomplete hematologic recovery (CRi), the LMPP/GMP compartments shrank in size (Figure 5C). Mature myeloid cells represented the greatest proportion of the BM and were either WT or contained the *ASXL1-*mutant clone which, by EOC1, had expanded 4-fold compared to baseline (Figure 5B). At EOC9, blood production was largely sustained by *ASXL1*-mutated and WT hematopoiesis that fully outcompeted leukemic clones within HSPCs (Figures 5B–5C). While *ASXL1* mutations confer unfavorable prognosis in AML,^18,20,49^ persistent *ASXL1* mutations at remission do not correlate with relapse in intensively treated patients.^50^ We now show that clones harboring a single *ASXL1* mutation can expand with IVO+VEN+AZA and do not necessarily lead to relapse.

**Figure 5.**
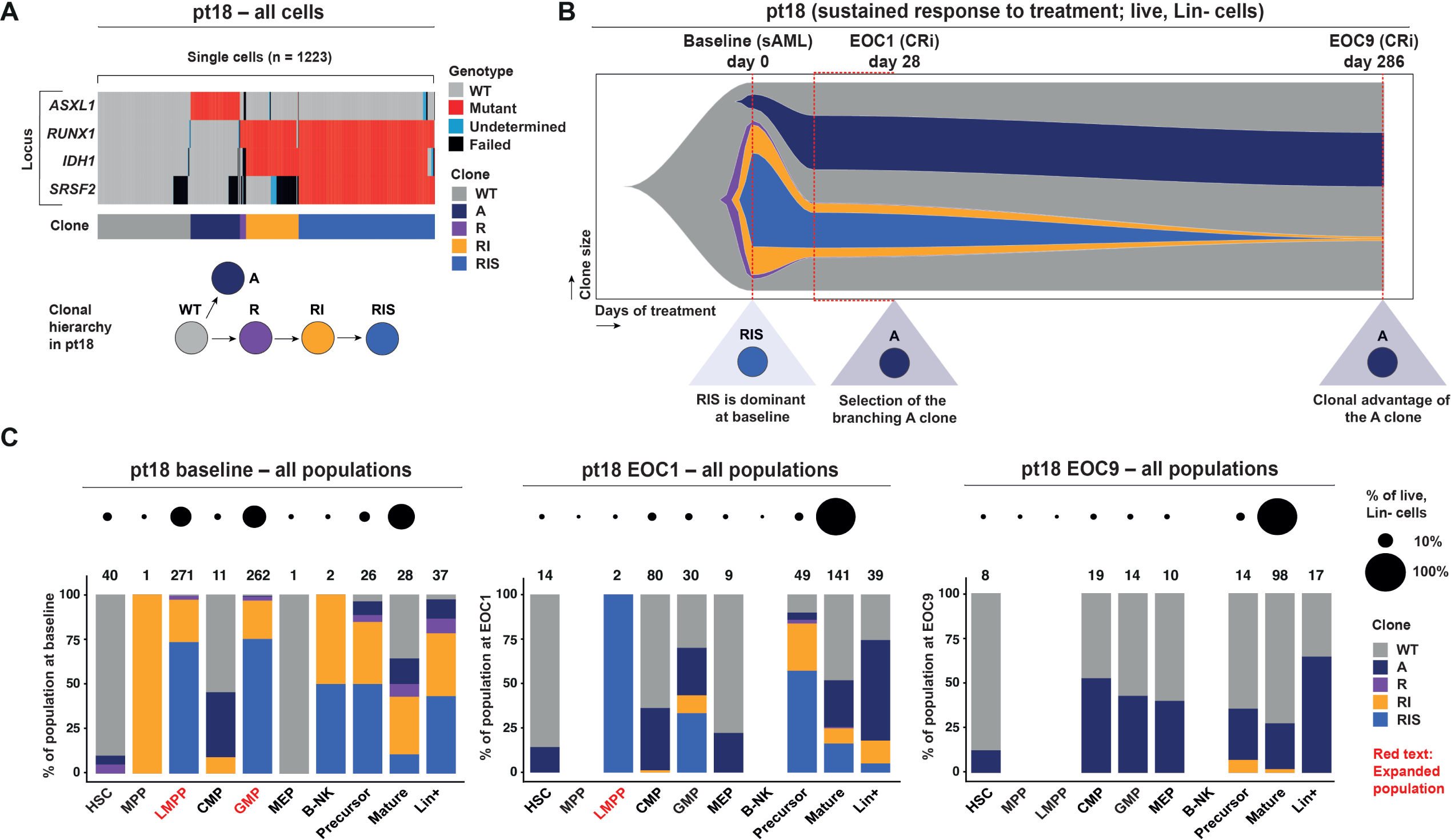
Selection of independent clonal hematopoiesis in the background of WT cells correlates with sustained response in pt18. (A) Same panel structure as in Figure 2A but for pt18 (n = 1223 cells). The genotyped mutations are: *ASXL1*-W583*, *RUNX1*-R204Q, *IDH1*-R132H, *SRSF2*-P95_R102del. Clone abbreviations are: WT: wild type (gray). A: *ASXL1* mutant (dark blue). R: *RUNX1* mutant (purple). RI: *RUNX1* and *IDH1* mutant (orange). RIS: *RUNX1*, *IDH1*, and *SRSF2* mutant (blue). (B) Same panel structure as in Figure 2B but for pt18, showing the baseline, EOC1, and EOC9 time points. Clone color-coding as in (A). sAML: secondary AML; CRi: complete remission with incomplete hematologic recovery. (C) Same panel structure as in Figure 2C but for pt18, across baseline, EOC1, and EOC9 time points. Clone color-coding as in (A).

Pt10 had been pretreated with 4 prior lines of therapy and relapsed after receiving allo-HSCT (Supplemental Figure 9). Remarkably, the dominant clone at baseline, containing 8 driver mutations, was fully eradicated by EOC3 (CRi; Supplemental Figures 9C–9E). WT donor hematopoiesis was reconstituted, with no evidence of selection of alternative leukemic clones (Supplemental Figure 9D; Supplemental Tables 1J and 4G). Pt10 has remained in remission after 57 months of follow-up. Here, in addition to eradicating leukemic/pre-leukemic clones, IVO+VEN may have given time for a donor graft versus leukemia effect. Notably, bulk NGS at remission revealed a *GNAS* mutation at 1% VAF, not detected pre-treatment (Supplemental Table 1H). Although not genotyped in TARGET-seq+, the mutation is likely to represent independent clonal hematopoiesis (CH).

Altogether, in patients who relapsed, resistant clones were selected within 3 treatment cycles, in progenitor or GM precursor compartments, with concurrent reduction of the dominant leukemic clones pre-treatment (Figure 6A). By contrast, in two patients who remained in sustained remission, therapy virtually eradicated leukemic clones from early compartments, allowing hemopoiesis from WT and CH cells.

**Figure 6.**
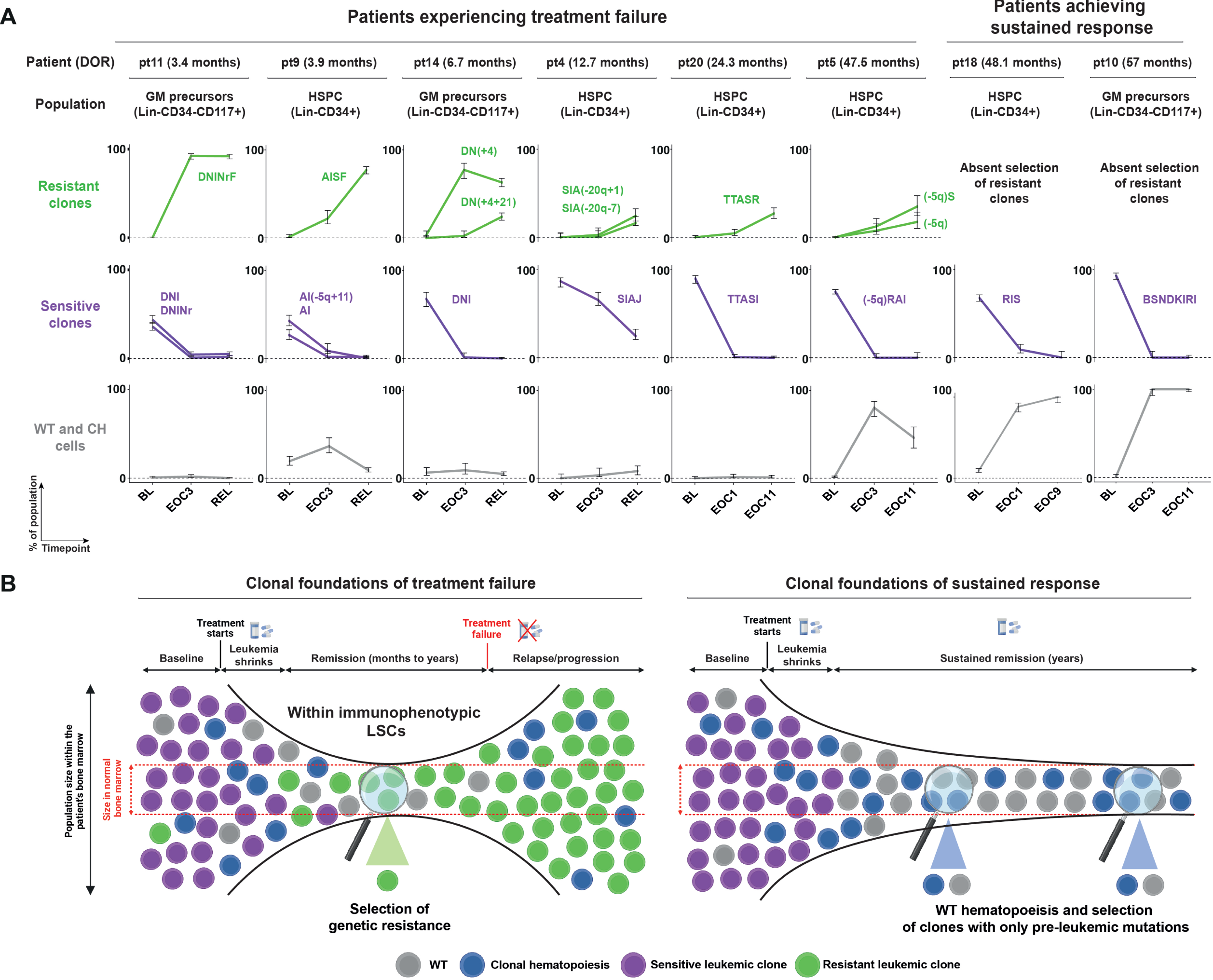
Rapid clonal selection within immature cell populations foreshadows patient outcome. (A) Line graphs showing longitudinal changes in clonal composition (indicated as % of all cells in a specific immunophenotypically-defined population, y-axis) in all studied patients (columns). The immunophenotypically-defined populations are indicated above. Clone abbreviations are indicated within the graphs. Clones associated with resistance are indicated in green; dominant pre-treatment clones sensitive to treatment are indicated in purple; the cumulative frequency of WT cells and clones harboring a single pre-leukemic driver mutation (i.e. CH clones) are indicated in grey. Dominant clones are defined as those clones contributing to at least 50% of the expanded immunophenotypic population, after arrangement in decreasing frequency (for multiple clones, the sum of their frequency was considered). Error bars depict the 95% confidence intervals calculated using the Wilson score interval.^59^ Time points are indicated on the x-axis. (B) Proposed model depicting the establishment of the clonal foundations of either treatment failure (left) or sustained response (right), early during treatment.

## Discussion

Despite the small size of our cohort and the remarkably complex variation of clonal dynamics across hematopoiesis under treatment, important general principles emerge. Therapy-driven clonal selection occurs rapidly, within 1-3 cycles of treatment, both in patients who relapse and those who remain in long-term remission (Figure 6B). Clonal selection leading to relapse initially occurred in small populations with LSC potential. Resistant clones were either not detected prior to therapy or were less fit pre-existing clones that expanded under therapeutic pressure and eradication of the primary leukemic clone. Rapid clonal selection within residual LSCs preceded treatment failure by months or even years; thus, the kinetics of initial selection did not correlate with the timing of frank hematological relapse. In contrast, in patients who remained in durable remission, only WT cells and CH were selected within HSPC, precursor, and mature myeloid compartments, raising the possibility of cure. Hence, we propose a model whereby rapid clonal selection by IVO+VEN±AZA within immature BM compartments early during treatment foreshadows longer-term patient outcome. Future studies need to address how genetic/epigenetic drivers cooperate with cell-specific programs within persistent LSCs to promote therapy resistance.

Our study has several implications. Persistent leukemic mutations at remission correlate with relapse in VEN+AZA^51,52^ and intensively treated patients.^50,53–55^ However, in addition to persistence of a leukemic mutation, we have identified that it is the early, rapid, and dynamic clonal re-organization within residual LSCs that builds the foundations of treatment failure. This extends our prior observations with enasidenib monotherapy.^1^ Furthermore, the temporal window between initial selection of relapse-associated clones and clinical relapse may provide an opportunity to therapeutically target mutant LSCs. These findings have important implications for MRD-directed therapies, as they become more available in the context of clinical trials, raising the possibility that selected clones may be eradicated prior to frank relapse. Finally, our findings emphasize that mechanistic investigation of clone-specific vulnerabilities should consider the differentiation stage at which clonal expansion occurs. For example, our observations in pt11 (Figure 2) suggest future studies should address how constitutive FLT3 signaling promotes therapy resistance specifically within LMPP/GMP/GM precursor populations. We and others have shown that the consequences of myeloid mutations vary between different stages of differentiation.^36–39^ Hence, it is plausible to speculate that the interaction between somatic genetics and cellular context may provide unique targetable vulnerabilities.

Despite rapid selection of resistant clones, the time interval between clonal selection and relapse varied from months to years. This may be due to: (a) variable clonal fitness; (b) different cells-of-origin of resistant clones; (c) clonal competition; (d) differing BM microenvironments. Though selection of pro-proliferative *FLT3*-ITD and *FLT3*-TKD mutations was associated with rapid relapse, not all clones bearing signaling mutations expanded at relapse (*NRAS* in pt9; *KRAS* and *PTPN11* in pt10; *BRAF* in pt20; and *CBL* in pt4); the clonal context or the variable proliferative properties of these mutations may have influenced their potency. Notably, clones bearing CNAs or a *RUNX1* mutation were also selected but associated with more delayed progression. Second-site *IDH1* mutations or *IDH2*-mutant clones were not detected, potentially secondary to the use of combined therapy as opposed to single agent IVO.

Interestingly, in pts 14/9/4, relapse-associated clones were already detected at baseline but only gained an advantage within LSCs under therapeutic pressure. The most parsimonious explanation is that therapy suppressed the dominant pre-treatment clones, enabling previously less fit clones to expand. Additionally, therapy may have induced non-genetic chromatin reprogramming,^56^ decreasing their differentiation capacity and/or enhancing self-renewal. Future linkage of single-cell clonal and chromatin readouts^37,57^ may identify putative non-genetic mechanisms of resistance. Finally, we cannot exclude that undetected/unknown coding or non-coding genetic drivers may have contributed to the expansion of pre-existing clones. Of note, even in cases where resistant clones were detected only after treatment exposure (pts 11/5/20), they may have been present already at diagnosis, albeit below the detection limit of conventional approaches, as previously suggested.^58^ Ultra-deep sequencing or ultra-sensitive targeted mutational profiling (applicable, for example, to mutations such as *FLT3*-ITD) at baseline may address this issue.

Collectively, our study highlights a common principle that clonal selection occurs early within populations containing LSC potential and foreshadows outcome to IVO+VEN±AZA. It will be important to determine if our findings extend to other targeted therapy combinations, as each treatment provides a selective pressure favoring the outgrowth of contextually fit clones.

## Authors’ Disclosures

**C.A. Lachowiez**: Consulting/Advisory: AbbVie, Servier, Rigel, Syndax, BMS, COTA healthcare. Research funding: AbbVie; **F. Ravandi**: Consulting/Advisory: BMS, Syros, AbbVie, Astellas, Prelude. Research funding: Astyex/Taiho, Syros, Amgen, Xencor, Prelude. **G. Issa**: Consulting/Advisory: Kura Oncology, Syndax, NuProbe, Novartis, AbbVie, Sanofi, AstraZeneca. Research funding: Celgene, Merck, Kura Oncology, Syndax, Astex, NuProbe, Novartis. **T. Kadia**: Consulting/Advisory: AbbVie, Genentech, BMS, Servier, Sellas, DrenBio. Honoraria: Novartis, Rigel. Research funding: AbbVie, Genentech, BMS, Jazz, Sellas, DrenBio, Regeneron, Amgen, Pfizer, Ascentage, Incyte, ASTEX, AstraZeneca, Cellenkos. **H. Kantarjian**: Consulting/Advisory/Honoraria: AbbVie, Amgen, Ascentage, Ipsen Biopharmaceuticals, KAHR Medical, Novartis, Pfizer, Shenzhen Target Rx, Stemline, Takeda. **M. Konopleva**: Consulting/Advisory: AbbVie, AstraZeneca, Auxenion GmbH, Bakx Therapeutics, Boehringer, Dark Blue Therapeutics, F. Hoffmann-LaRoche, Genentech, Gilead, Immune Oncology, Janssen, Legend Biotech, MEI Pharma, Redona, Sanofi Aventis, Sellas, Menarini Group, Vincerx. Research funding: AbbVie, Allogene, AstraZeneca, Genentech, Gilead, ImmunoGen, MEI Pharma, Precision Biosciences, Rafael Pharmaceutical, Sanofi Aventis, Menarini Group. Clinical trials: AbbVie, AstraZeneca, Cellectis, Genentech, Janssen, Pfizer, Sanofi Aventis, Menarini Group. IP: Reata Pharmaceutical. **C.D. DiNardo**: Consulting/Advisory/Honoraria: AbbVie, AstraZeneca, BMS, Genentech, GenMab, GSK, Immunogen, Notable Labs, Rigel, Schrodinger, Servier, Amgen, Astellas, Gilead, Jazz, Stemline; Research Funding: AbbVie, BMS, Servier, Astex, ImmuneOnc, Cleave, Foghorn, Loxo, Rigel. **P. Vyas**: Consulting/Advisory: AbbVie, Servier, Rigel, Syndax, AstraZeneca, Debiopharm, Charm Therapeutics. Research funding: Bristol Myers Squibb. Co-founder and Board: Yellowstone Biosciences. SAB: Auron Therapeutics. No disclosures were reported by the other authors.

## Authors’ Contributions

**S. Turkalj:** Data curation, formal analysis, investigation, methodology, visualization, writing-original draft, writing-review & editing. **B. Stoilova:** Conceptualization, data curation, formal analysis, investigation, methodology, writing-review & editing. **A. Groom:** Investigation, methodology, writing-original draft, writing-review & editing. **F.A. Radtke:** Formal analysis, methodology, visualization, software, writing-original draft, writing-review & editing. **R. Mecklenbrauck:** Data curation, visualization, writing-original draft, writing-review & editing. **N.A. Jakobsen:** Investigation, methodology, software, writing-review & editing. **C.A. Lachowiez:** Data curation, writing-review & editing. **M. Metzner:** Data curation, formal analysis, methodology, writing-review & editing. **B. Usukhbayar:** Methodology, writing-review & editing. **M.A. Salazar:** Methodology, writing-review & editing. **Z. Zeng:** writing-review & editing. **S. Loghavi:** Data curation, writing-review & editing. **J. Marvin-Peek:** Data curation, writing-review & editing. **V. Körber:** Formal analysis, visualization, writing-review & editing. **F. Ravandi:** Writing-review & editing. **G. Issa:** Writing-review & editing. **T. Kadia:** Writing-review & editing. **V. Symeonidou:** Methodology, writing-review & editing. **A.P. de Groot:** Methodology, writing-review & editing. **H. Kantarjian:** Writing-review & editing. **K. Takahashi:** Resources, writing-review & editing. **M. Konopleva:** Conceptualization, resources, writing-review & editing. **C.D. DiNardo:** Conceptualization, resources, writing-review & editing. **P. Vyas:** Conceptualization, resources, funding acquisition, supervision, project administration, writing-original draft, writing-review & editing.

## Acknowledgements

P.V. acknowledges funding from the Medical Research Council (MRC) Molecular Haematology Unit Programme Grant (MC_UU_00029/8), Blood Cancer UK Programme Continuity Grant 13008, NIHR Senior Fellowship and the Oxford BRC Haematology Theme. P.V. and B.S also acknowledge previous funding from the MRC Molecular Haematology Unit Programme Grant (MC_UU_12009/11). S.T. is supported by the WIMM DPhil Prize Studentship funding from Scatcherd European Scholarship in partnership with The Medical Research Council/Radcliffe Department of Medicine and The Clarendon Fund. A.J.G. is supported by DPhil studentship funding from the Lady Tata Memorial Trust. F.A.R. is supported by the Oxford – Sir David Weatherall Scholarship from Green Templeton College and the WIMM Prize Studentship from the MRC Weatherall Institute of Molecular Medicine. R.M. is supported by the Mildred-Scheel-Postdoc Program by the Deutsche Krebshilfe (project number 70115737). C.D.D is supported by the LLS Scholar in Clinical Research Award. N.A.J. was supported by a Medical Research Council and Leukaemia UK Clinical Research Training Fellowship (MR/R002258/1). M.M., B.U., and M.A.S. were funded by the Haematology Theme of the Oxford NIHR Biomedical Research Centre. V.K. was funded by the Deutsche Forschungsgemeinschaft (DFG, German Research Foundation – project number 526169089). This research was funded in part by the NIH/NCI Cancer Center Support Grant P30 CA016672. The authors thank Prof. James Davies, Prof. Peter Valk, Prof. Michael Heuser, Dr. Susann Rahmig, Dr. David Cruz Hernandez, and Ms. Grace Meaker, for insightful comments and discussions. The authors also acknowledge the MRC WIMM Flow Cytometry and Single Cell Facilities. Biorender was used fully or in part to generate the following figures: Figures 1B and 6B; and Supplemental Figures 4C, 6E, 7F, 8A, and 9D.

## Notes

### Competing Interest Statement

The authors have declared no competing interest.

