## Supplemental Materials and Methods for "Rapid clonal selection within early hematopoietic cell compartments presages outcome to ivosidenib combination therapy"

#### Supplemental Methods

##### Bulk DNA sequencing

For the 81-gene targeted NGS panel, interrogating 81 genes recurrently mutated in myeloid malignancies, genomic DNA (gDNA) was extracted from the buffy coat of fresh BM aspirate samples. Libraries were generated by Haloplex probe capture followed by PCR amplification of target regions using Haloplex chemistry (Agilent, Santa Clara, CA). Sequencing was performed on the MiSeq (Illumina, San Diego, CA) using paired-end reads. Human genome build 19 (hg19) was used as the reference for read alignment. Agilent SureCall was used for variant calling. Variant allelic frequency was estimated as the percentage of variant reads over total reads at a specific locus. Matched germline DNA was not available for comparison. Sensitivity was established at 1–2%.

For the 97-gene targeted NGS panel, interrogating 97 genes recurrently mutated in myeloid malignancies, pre-capture DNA libraries were prepared using the KAPA HyperPlus protocol (Roche). DNA was fragmented by enzymatic fragmentation, followed by end repair and A-tailing, after which adapter ligation was performed using KAPA dual-indexed adapters (Roche). Library cleanup was performed using Agencourt AMPure XP beads. Amplification was performed by ligation-mediated PCR using a KAPA HiFi HotStart high-fidelity DNA polymerase (Roche), followed by AMPure XP bead purification. Targeted capture was performed using a custom pool of biotinylated capture probes (SeqCap EZ Prime Choice, Roche) spanning 347 kb (Supplemental Table 1). Amplified DNA libraries were hybridized to the capture probes. The captured libraries were amplified by 14 cycles of PCR using a KAPA HiFi HotStart DNA polymerase and purified using AMPure XP Beads. Post-capture libraries were pooled in equimolar concentrations and were sequenced on an Illumina NextSeq 500 using paired-end reads with an average depth of 822x.

WES analysis for mutation discovery was performed using $CD45^{dim}$ blasts and $CD3^{+}$ T control cells were sorted on a BD Fusion machine. gDNA was extracted using DNA Mini or Micro Kits (Qiagen). WES was performed by Novogene Europe using low input library preparation method and SureSelect Human All Exon V6 bait design (Agilent). Libraries were sequenced using PE150 reads 50x for T cells and 100x for blasts. Data was mapped using bwa (bwa 0.7.17). Variant calling was performed using Mutect2 (gatk 4.0.5.1), Strelka2 (strelka 2.9.2) and vardict (vardict=2018.10.18) and taking the intersect of two out of the three callers and VAF > 0.05. *FLT3*-ITD was additionally called using Pindel (pindel 0.2.5).

##### Flow cytometry and FACS

Patient samples were first thawed at 37°C in a water bath, followed by addition of warm fetal bovine serum (FBS) and thawing media (IMDM supplemented with 20% FBS and 110 µg/mL

DNase). Cells were centrifuged, resuspended in FACS staining buffer (IMDM with 10% FBS and 10 µg/mL DNase), placed on ice, counted, and stained with the following antibodies: anti-CD38-BV421 (1:20, Biolegend, clone HIT2), anti-CD10-BV605 (1:40, Biolegend, clone HI10a), anti-CD117-PE/Dazzle594 (1:80, Biolegend, clone 104D2), anti-CD45RA-BB515 (1:40, BD, clone HI100), anti-CD123-PE (1:40, Biolegend, clone 6H6), anti-CD90-PE/Cy7 (1:20, Biolegend, clone HI100), anti-CD34-APC (1:160, Biolegend, clone 581), using a lineage cocktail containing: anti-CD2-PE/Cy5 (1:160, Biolegend, clone RPA-2.10), anti-CD3-PE/Cy5 (1:320, Biolegend, clone HIT3a), anti-CD4-PE/Cy5 (1:160, Biolegend, clone RPA-T4), anti-CD8a-PE/Cy5 (1:320, Biolegend, clone RPA-T8), anti-CD19-PE/Cy5 (1:160, Biolegend, clone HIB19), anti-CD20-PE/Cy5 (1:160, Biolegend, clone 2H7), anti-CD235ab-PE/Cy5 (1:320, Biolegend, clone HIR2). Cells were incubated for 30–45 min on ice, washed with 1 mL of FACS staining buffer, centrifuged, and resuspended in FACS staining buffer with 7-AAD (Biolegend) live/dead stain at 1:100 dilution.

Flow cytometry analysis and sorting for TARGET-seq+ were performed on a Sony MA900 sorter. For each patient, samples from different time points were processed in the same sort (except for pt5, where the EOC11 sample was processed and sorted separately from baseline and EOC3 samples). Unstained, single-stained, and fluorescence-minus-one (FMO) controls were used. FACS sorting gates were set using a combination of FMO controls and a stained healthy WT BM sample (NOC156), in which population sizes were known *a priori*. For samples with a considerable amount of Lin$^+$ cells, Live/Lin$^-$CD34$^+$, Live/Lin$^-$CD34$^-$CD117$^+$, and Live/Lin$^-$CD34$^-$CD117$^-$ cells were first pre-sorted. Single cells were then index-sorted into 384-well plates containing 3 µL TARGET-seq+ lysis buffer (below). Doublets and dead cells were excluded. The following populations were sorted: Live/Lin$^-$CD34$^+$, Live/Lin$^-$CD34$^-$CD117$^+$, and Live/Lin$^-$CD34$^-$CD117$^-$ cells. Additionally, for pts 4/9/11/14, specific HSPC subpopulations within Live/Lin$^-$CD34$^+$ cells were enriched. Cells from the NOC156 control sample were sorted onto every plate, making up ~10% of wells. Plates were centrifuged and snap frozen on dry ice prior to storage at –80°C.

**TARGET-seq+ library preparation**

***Primer design***

All loci were genotyped using primers annealing either to the introns flanking the mutation, or to the exon containing the mutation, which allowed genotyping from gDNA. Additionally, where possible, cDNA primers were designed to anneal to exons outside of the mutant exon, so that genotyping reads derived from cDNA could be obtained as well. Primer-BLAST was used for primer design. The specificity and efficiency of each pre-amplification and nested primer pair were tested first in bulk PCR reactions and then in single human CD34$^+$ BM cells.

***TARGET-seq+***

The oligo(dT)-lysis buffer mix was prepared containing 0.1% Triton X-100 (Sigma-Aldrich), 0.5 mM dNTPs (Life Technologies), 5% PEG 8000 (Sigma-Aldrich), 0.5 U/µL RNase inhibitor (Takara), 2.7 x $10^{-05}$ AU/µL protease (Qiagen), 1:8,000,000 diluted ERCC spike-in mix (Ambion), and 668 nM of barcoded oligo(dT)-ISPCR primer. 3 µL of barcoded oligo(dT)-lysis buffer mix were transferred into each well of 384-well plates. Single cells were index-sorted as described above. Plates were incubated at 72°C for 15 min followed by the addition of 1 µL of the reverse transcription (RT) mix, bringing the reaction to 25 mM Tris-HCl (Thermo Scientific), 30 mM NaCl (Invitrogen), 2.5 mM $MgCl_2$ (Invitrogen), 1 mM GTP (Thermo Scientific), 8 mM Dithiothreitol (DTT, Thermo Scientific), 0.5 U/µl RNase inhibitor (Takara), 2 µM of Smart-seq2 template switching oligo (TSO, IDT), 2 U/µl of Maxima H-minus reverse transcriptase enzyme (Thermo Scientific), and 70 nM cDNA primers. RT was carried out at 42 °C for 90 min, followed by 10 cycles of 50 °C for 2 min and 42 °C for 2 min, with a final incubation at 85 °C for 5 min. 6 µL of PCR mix were then added to achieve 1 × KAPA HiFi HotStart Ready Mix (Roche), 50 nM ISPCR primer, 400 nM genotyping gDNA primers, and, where applicable, 28 nM genotyping cDNA primers. The following PCR program was used: 98 °C for 3 min, 21 cycles of 98 °C for 20 s, 67 °C for 30 s, and 72 °C for 6 min, followed by incubation at 72 °C for 5 min. The resulting cDNA/genotyping library was split into two aliquots: one was used for 3' biased whole transcriptome library construction while the other for single-cell genotyping library construction.

For 3' biased transcriptome library construction, 1 µL of cDNA/genotyping library was pooled from each well, keeping each plate separate. Pools were purified twice using Ampure XP beads with 0.6:1 beads to cDNA ratio, quantified, and checked using Bioanalyzer (Agilent). A Nextera XT DNA Library Preparation Kit (Illumina) was used for tagmentation-based library preparation, using custom PCR primers (Supplemental Table 3). 4 ng from each pool (5 µL) was mixed to 5 µL of ATM enzyme and 10 µL of TD buffer. The reaction was incubated at 55 °C for 10 min and stopped with 5 µL of neutralization buffer (Illumina). 5 µL Nextera XT i7 forward index primer (Illumina), 5 µL custom i5 index primers (2 µM; sequences in Supplemental Table 3), and 15 µL of NPM enzyme were added and the reaction incubated with the following program: 95 °C for 30 s, 14 cycles of 95 °C for 10 s, 55 °C for 30 s and 72 °C for 30 s, and a final incubation at 72 °C for 5 min. Libraries were purified twice with Ampure XP beads using a 0.7:1 beads to cDNA ratio, quantified, and checked using Bioanalyzer. Libraries were pooled at equimolar ratios and sequenced with custom primers for Read1 and Index2 (Supplemental Table 3). Libraries for samples from all patients except pt5 were sequenced on a NovaSeq S4 flow cell (Illumina) with a targeted sequencing depth of 1 million reads/cell and the following configuration: 15 bp R1; 8 bp index read 1; 8 bp index read 2; 200

bp R2. Samples from pt5 were sequenced on a NextSeq Mid Output v2.5 Flow Cell (Illumina) with a targeted sequencing depth of 40,000 reads/cell and the following configuration: 15 bp R1; 8 bp index read 1; 8 bp index read 2; 69 bp R2.

To generate single-cell genotyping libraries, two PCR steps (PCR1 and PCR2) were performed. Nested genotyping primer sequences and barcodes used for PCR1 are listed in Supplemental Table 3. PCR1 was performed using 3.25 µL of KAPA 2G Robust HS Ready Mix (Sigma-Aldrich), 1.5 µL of cDNA/genotyping library, and 300 nM nested genotyping primers. PCR2 was performed using FastStart High Fidelity polymerase (Sigma-Aldrich) with 1.0 µL of PCR1 product and 1.2 µL of each barcoded primer (Access Array Barcode Library for Illumina Sequencers - 384, Single Direction, Fluidigm). 1 µL of the resulting fully indexed amplicons were pooled from each well, keeping plates separate, and purified with Ampure XP beads using a 0.8:1 beads to PCR product ratio. Libraries were quantified and checked with Tapestation (Agilent). Libraries were pooled at equimolar ratios, the final pool diluted to 10 pM in HT1 buffer, and sequenced on a NextSeq 500/550 Mid Output v2.5 kit (300 cycles) (Illumina), using 150 bp paired-end reads, with 10 bp for the index read. Custom sequencing primers were used (Supplemental Table 3).

**Targeted single-cell genotyping analysis**

***Single-cell genotyping pre-processing***

For amplicons containing single nucleotide changes (SNVs), the TARGET-seq pipeline (https://github.com/albarmeira/TARGET-seq) was used for pre-processing of single-cell genotyping reads. Reads were demultiplexed using barcodes introduced during PCR1 and PCR2, to generate cell-specific FASTQ files. For amplicons containing single nucleotide variants (SNVs), reads were aligned to human genome build 38 (hg38) using STAR version 2.7.3a with default settings. Where applicable, cDNA and gDNA amplicons were separated, extracting reads matching the gDNA and cDNA nested primer sequences. *mpileup* (samtools version 1.1) was used for variant calling, using --*minBQ 30*, --*count-orphans*, --*ignore overlaps*.

For amplicons containing small indels, WT and indel reads were counted from single-cell FASTQ files using the *grep* function in Unix. If the indel-containing locus was covered by both read 1 and read 2, counting was performed from both on R1 and R2 FASTQ files, providing the reverse complement sequences if counting was done from R2 files. The total WT and indel read count were then calculated.

In the specific case of *FLT3*-ITD (pt11), the getITD pipeline, adapted for single-cell analysis, was used to count the number of ITD and WT reads from each single cell.

##### ***Mutation calling***

Single-cell coverage was computed across the mutant locus. For most loci, cells with coverage < 30 reads were excluded (i.e. the amplicon is called undetected due to failed single-cell genotyping). For a small number of loci, this threshold was relaxed due to low coverage across most single cells. Specifically, this applies to the gDNA amplicon for *NPM1* in pt11 and pt14 (threshold: 15 reads); gDNA and cDNA amplicons for *DNMT3A* in pt11 (threshold: 5 reads); and the gDNA amplicon for *BRINP3* in pt10 (threshold: 15 reads). The single-cell variant allele frequency (scVAF) for each locus was calculated as:

*scVAF = variant reads / coverage*

To define scVAF thresholds for mutation calling at each locus, single cells from a control BM sample that was WT for all mutations of interest (from sample NOC156) were sorted onto every plate and processed simultaneously with patient cells. scVAFs at each locus were defined as stated above. The resulting scVAF distribution from WT control cells was used to define thresholds for genotype assignment at each locus. The scVAF threshold for calling a cell WT was set as:

*scVAF threshold for calling a cell WT = mean(scVAF in WT control cells) + 3 × SD of scVAF in WT control cells*

If the scVAF was below this threshold, the cell was called WT. The scVAF threshold for calling a cell mutant was then set as:

*scVAF threshold for calling a cell mutant = mean(scVAF in WT control cells) + 3 × SD of scVAF in WT control cells + 0.01*

If the scVAF was above this threshold, and if the cell yielded at least 10 mutant reads, the cell was called mutant (the minimum mutant read number was relaxed for the few exception cases noted). Cells with a scVAF between the two thresholds, or where the number of mutant reads was < 10, were called undetermined. In rare instances, a WT control cell showed an aberrantly high scVAF (> 0.1), due to cross-well contamination. In those cases, these cells were eliminated from threshold calculation, as their inclusion would skew the threshold value, causing inaccurate genotyping. Numbers of control cells eliminated were 2 for pt18; 1 for pts 4/10; and 8 for pt9. If cDNA reads were also available, the above strategy was performed separately for gDNA and cDNA amplicons. A consensus single-cell genotype was then assigned as follows: If the mutation was detected from either gDNA or cDNA, the cell was called mutant; if the gDNA amplicon was WT and the cDNA amplicon was not mutant, the cell was called WT; if the gDNA amplicon failed single-cell genotyping and the cDNA amplicon was WT, the cell was called undetermined.

##### *Single-cell transcriptome data pre-processing*

Transcriptome sequencing reads were demultiplexed via *bcl2fastq* using unique i7-i5 index combinations. Reads were then further demultiplexed using 14 bp cell barcodes using cutadapt v3.4. cDNA reads were simultaneously trimmed for polyA tails, Nextera adapters, and low-quality reads. Reads were mapped to the hg38 reference genome with STARsolo v2.7.10a, using the GENCODE v38 reference gene annotation, which included only protein-coding genes and long non-coding RNAs. Counts for each gene were obtained using default parameters and *--soloType SmartSeq --soloFeatures GeneFull_Ex50pAS*. Quality metrics were calculated with FastQC (v0.11.9), Samtools flagstat (v1.12), MultiQC (v1.11), and STAR outputs.

Single-cell transcriptome pre-processing was performed using Seurat (v5.0.1) and SingleCellExperiment (v1.24.0). Metadata with single-cell genotyping and FACS index data were first matched with single-cell identifiers based on the plate and well coordinates. Cells meeting the following filtering criteria were included: reads assigned to genes ≥ 25,000; genes detected ≥ 2,000; reads assigned to ERCC transcripts ≤ 50%; reads in mitochondrial genes ≤ 15%. A Seurat object was then generated with raw counts, excluding those cells that did not pass QC.

##### *inferCNV analysis and CNA calling*

To identify CNAs at single-cell level from scRNA expression data in pts 4/5/9/14, inferCNV was used (v1.18.1, https://github.com/broadinstitute/infercnv). First, for each patient, the *CreateInfercnvObject* command was runn, providing the patient-specific counts matrix from the Seurat object, which contained all the cells from that patient plus all healthy control BM cells passing scRNA-seq QC.

To enable CNA calling for samples from pt5, run on lower sequencing depth compared to other patients (counts on average 14,369 versus 1,919,130, respectively), probabilistic downsampling of the healthy control BM cells was performed. A scaling factor was calculated by dividing the average number of counts across cells from patient 5 and healthy control BM cells sequenced with patient 5 by the average number of counts across cells from healthy control BM samples which were sequenced to a higher depth. A distribution of probabilities was calculated by dividing the counts for a given gene by the total number of counts. These probabilities were then used to weigh the random selection of counts to remove in order to maintain an accurate representation of count distributions for the genes.

To identify CNAs with the HMM algorithm, the *infercnv* command was run using the following parameters: cutoff = 1, denoise = TRUE, HMM = TRUE, analysis_mode = “subclusters”, hclust_method = “ward.D2”, leiden_resolution = 0.01, tumor_subcluster_pval = 0.05. To identify high-confidence CNAs, only those regions previously detected by karyotyping (Supplemental Table 1) were taken forward for single-cell analysis. In addition, specifically in pt4, the loss on chromosome 7 was included, which showed a strong CNA signal in a subset of cells but was not detected in karyotyping data. The set of regions containing *bona fide* CNAs obtained by this approach was subjected to single-cell analysis.

Specifically, for each single cell, it was calculated whether the average gene expression in the region of interest (either the entire chromosome for very large CNAs, or a manually selected portion of the chromosome for shorter CNAs) was significantly higher or lower than expected in a diploid cell. The standard deviation of the average read count per cell and gene across the entire genome (which, on average, is diploid in our samples) was computed as

$$\sigma_{\langle n\rangle}=\sqrt{G}\sqrt{\frac{1}{K}\sum_{k=1}^{K}\left(\langle n\rangle_{G,k}-\mu_{\langle n\rangle_G}\right)^2},$$

where $G$ is the total number of genes that went into the analysis, $K$ is the total number of cells (both taken from the final infercnv object), $\langle n\rangle_{G,k}$ is the average number of normalized read counts per gene in cell $k$, and $\mu_{\langle n\rangle_G}$ is the average of $\langle n\rangle_G$ across all cells (both estimated from all $G$ genes). Single cells were then classified as affected by a particular CNA if the gene expression in the respective region was significantly higher (in the case of copy number gains) or lower (in the case of copy number losses) than expected by chance. To this end, the average gene expression per cell was computed on the subset of genes, $g$, that is affected by a particular CNA of interest (specified in the respective figure legends), yielding an estimate $\langle n\rangle_{g,k}$ for each cell $k$. Accepting a false discovery rate of 5% and assuming that average read counts per cell are normally distributed, cells were classified as harboring a copy number gain if $\langle n\rangle_{g,k} > \mu_{\langle n\rangle_g} + 1.96\frac{\sigma_{\langle n\rangle}}{\sqrt{g}}$, as harboring a copy number loss if $\langle n\rangle_{g,k} < \mu_{\langle n\rangle_g} - 1.96\frac{\sigma_{\langle n\rangle}}{\sqrt{g}}$, and else as diploid. For pt4, pt9, and pt14, CNA calls in single cells were then merged with SNV calls for the same cells, and infSCITE was performed on integrated SNV+CNA calls to establish the clonal hierarchy (below).

***Inference of clonal hierarchies***

Once mutations and, where applicable, CNAs, were called at all loci in a single patient, the pattern of mutational co-occurrence was used to determine clonal structures and assign cells to clones. For pts 10/11/18/20, genotyping calls for SNVs and small indels were used. For pts 4/5/9/14, genotyping calls for SNVs, small indels, and CNAs (as determined by inferCNV) were used.

First, mutation calls were integrated across all cells from a given patient, for all genotyped loci. infSCITE was used to determine the statistically most likely phylogenetic tree. As input, the matrix containing the mutational status for each locus in each cell was used. infSCITE was run with default parameters and '-r 200 -L 10000 -fd 0.01 -ad 0.02 -e 0.2 -p 1000'. The output phylogenetic tree informed the clonal assignments for single cells. We first checked that each phylogenetic tree was broadly consistent with VAFs obtained by bulk sequencing. In two specific cases where we could not confidently assign the exact course of mutation acquisition (pts 10 and 11), this was stated in the respective figure legends. In pt11, a substantial number of cells failed single-cell genotyping for *DNMT3A*-N757K locus (Figure 2A) and infSCITE predicted *DNMT3A*-N757K to be subclonal to *NPM1*-W288fs*12. However, prior literature suggested this to be an unlikely course of events. WES indicated a VAF of 47% for *NPM1*-W288fs*12 and 44% for *DNMT3A*-N757K, suggesting both mutations were early events. In the specific case of pt10, the infSCITE output was unreliable due to a high predicted "ad" value. In the case of pt4, further manual curation of the phylogenetic trees was performed. In this case, inferCNV and SNV analysis suggested that gain of chromosome 1 (chr1+) occurred twice, within independent SIA(+20) and SIAJ clones. Due to chr1+ being treated as a single event in the infSCITE input, these two clones were manually assigned.

Due to allelic drop-out (ADO), a mutation is occasionally not detected in a subset of mutant cells. When ADO occurred in a locus harboring an early mutation, but when later somatic mutations were detected in the same cell, that cell could still be assigned to the right daughter clone, (i.e. the later mutation acted as a 'control' event). For example, in pt18, there were 14 cells mutant for *SRSF2* but WT for *IDH1* (Figure 5A); given that the *IDH1* mutation occurred prior to the *SRSF2* mutation, these cells were still assigned to the *SRSF2* mutant, *IDH1* mutant RIS clone. In cases where a branching clonal structure was present, a small proportion of cells contained mutations which were mutually exclusive as per clonal structure (either due to a doublet or due to cross-well contamination). In these cases, the clone was not assigned.

**Analysis of FACS index data**

Flow cytometry index data were recorded for each single cell during single-cell sorting. Fluorescence values were recorded for forward scatter (FSC), back-scatter (BSC; equivalent to side scatter, SSC), and all fluorochromes used. Index data were matched with single-cell identifiers based on the well barcodes and combined with single-cell clonal calls. Cells were labeled as positive or negative for each surface marker, based on FACS gating, and assigned to immunophenotypic populations. For example, cells that were $Lin^{-}CD34^{+}CD38^{-}CD10^{-}$ $CD45RA^{-}CD90^{+}$ were labeled as immunophenotypic HSC.

**Inference of clone size within Lineage-negative cells**

For pts 4/5/9/10/11/14/18, the size of each clone within all live, Lineage-negative ($Lin^-$) cells at each time point was inferred. For each sample, we first computed the fraction of each clone within each immunophenotypic population, and the upper and lower limits of the 95% confidence interval (CI) were calculated via the Wilson score interval. We then multiplied both the population-specific clonal fractions and the upper and lower limits of the CI with the relative size of the respective immunophenotypic population (measured by flow cytometry as the fraction of that population within all sampled $Lin^-$ populations). The obtained population-specific clonal fractions were summed for each clone, to infer the total size of each clone within all sampled $Lin^-$ populations. The CI limits (multiplied with the relative size of the respective immunophenotypic population) were also summed for each clone, to compute the 95% CI of the inferred clone size within $Lin^-$ cells. The results for each patient are summarized in Supplemental Table 4. In all cases in which one or several $Lin^-$ were not sampled, those populations contributed cumulatively to < 1% of all $Lin^-$ cells. This analysis was not done for pt20, in which < 5 cells were sampled from the largest $Lin^-$ population at EOC1 (mature myeloid population), thus precluding high-confidence inference of clone size within all $Lin^-$ cells.

### Supplemental Figures and Legends

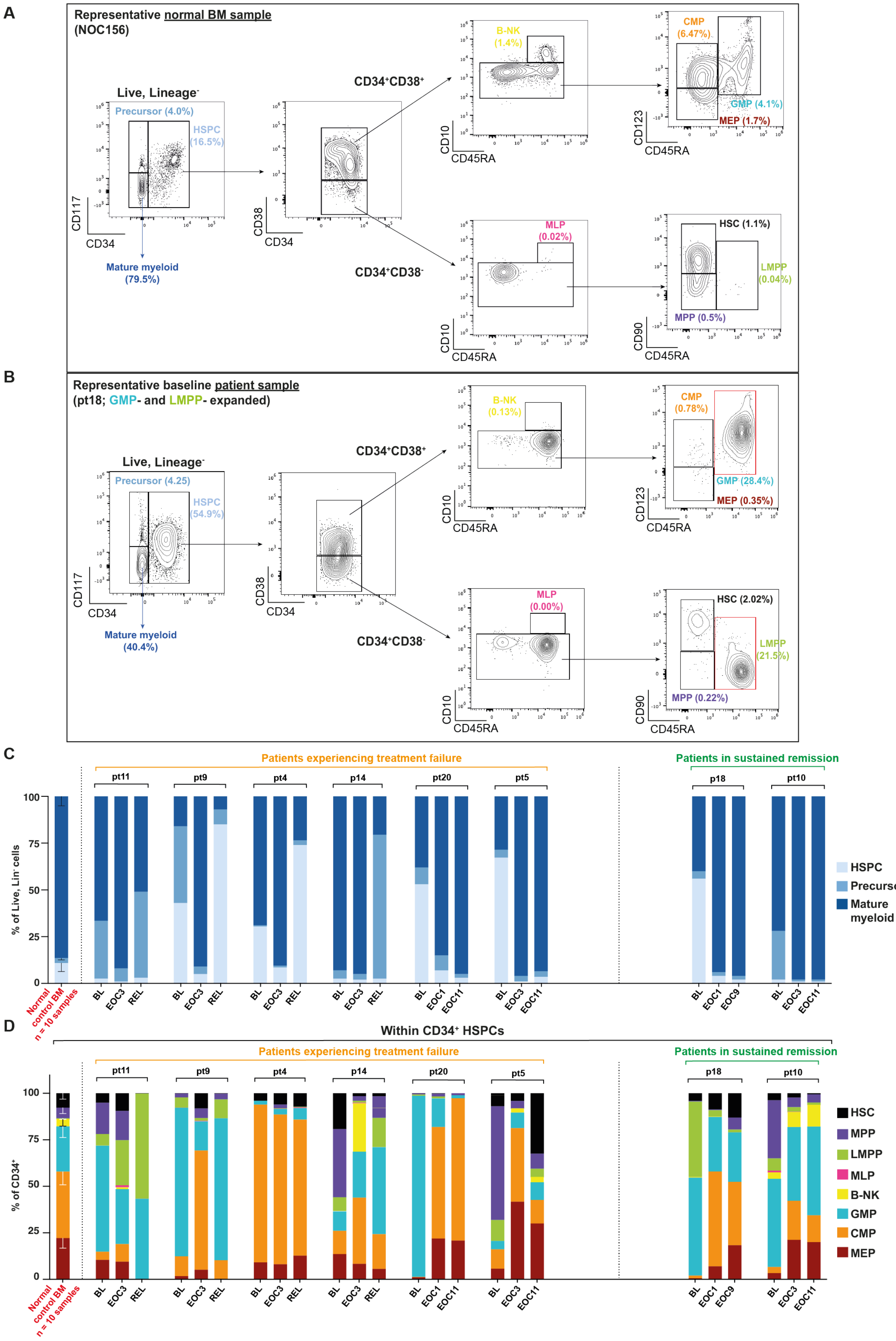

**Supplemental Figure 1. Flow cytometry panel and immunophenotypic analysis of all patients studied.**

(A) Representative flow cytometry panel (the normal NOC156 bone marrow sample shown) used for single-cell index sorting and flow cytometry analysis. Size of each population as percentage of all live, $Lin^-$ cells is indicated. HSPC: hematopoietic stem and progenitor cells ($Lin^-CD34^+$); HSC: hematopoietic stem cell ($Lin^-CD34^+CD38^-CD10^-CD90^+CD45RA^-$); MPP: multipotent progenitor ($Lin^-CD34^+CD38^-CD10^-CD90^-CD45RA^-$); LMPP: lymphoid-primed multipotent progenitor ($Lin^-CD34^+CD38^-CD10^-CD90^-CD45RA^+$); CMP: common myeloid progenitor ($Lin^-CD34^+CD38^+CD10^-CD123^{mid}CD45RA^-$); GMP: granulocyte-monocyte progenitor ($Lin^-CD34^+CD38^+CD10^-CD123^{mid/hi}CD45RA^+$); MEP: megakaryocyte-erythroid progenitor ($Lin^-CD34^+CD38^+CD10^-CD123^-CD45RA^-$); MLP: multi-lymphoid progenitor ($Lin^-CD34^+CD38^-CD10^+$); B-NK: B-cell/NK-cell progenitor ($Lin^-CD34^+CD38^+CD10^+$).

(B) As in (A) but showing the baseline sample from pt18. The red gates indicate aberrantly expanded populations (LMPP and GMP in this case).

(C) Immunophenotyping of HSPC ($Lin^-CD34^+$), GM precursor ($Lin^-CD34^-CD117^+$), and mature myeloid ($Lin^-CD34^-CD117^-$) populations in patient samples (x-axis; BL, baseline; EOC, end of cycle; REL, relapse). Populations are quantified as percentage of all live, $Lin^-$ cells (y-axis). Color legend on the right. Patient identifiers shown above. As control, healthy BM (n = 10 biologically independent samples) was used (leftmost bar). Error bars indicate the standard deviation (sd) of the mean.

(D) Detailed composition of the HSPC sub-compartments in patient samples and healthy BM controls (x-axis). HSPC sub-populations are quantified as percentage of all $Lin^-CD34^+$ cells (y-axis). Color legend on the right. The rest as in (C).

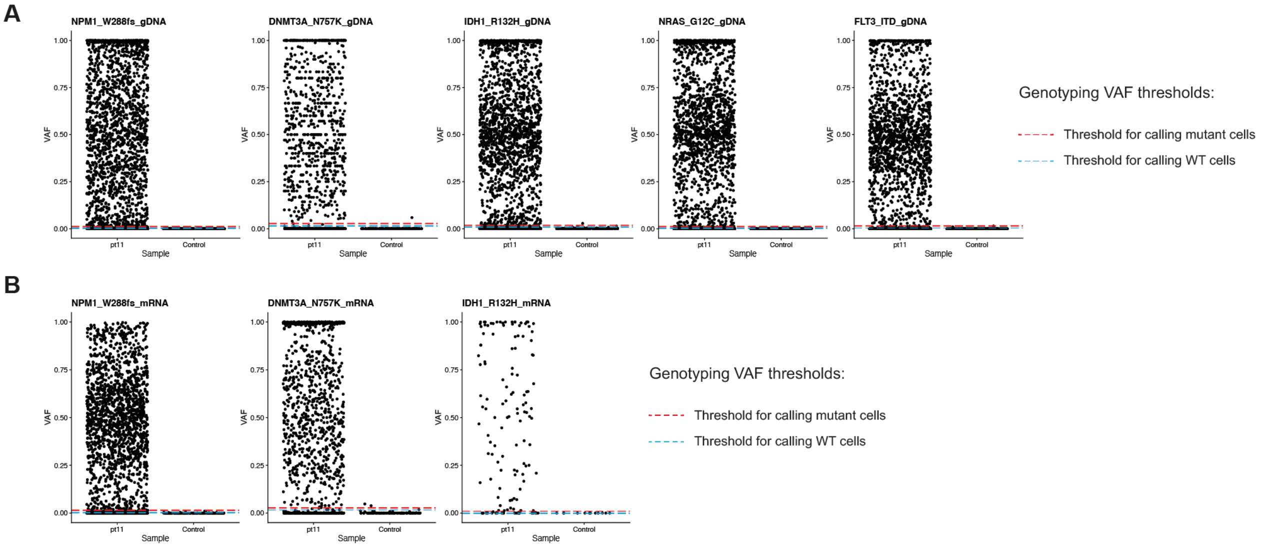


**Supplemental Figure 2. Single-cell genotyping for pt11.**

(A) Single-cell variant allele frequency (scVAF) distribution (y-axis, linear scale) from genomic DNA (gDNA) amplicons across five mutant loci in pt11 genotyped by TS+. scVAFs from pt11 are shown on the left, and scVAFs from a healthy control BM wild-type (WT) for all genotyped loci are shown on the right of each graph. The mutations are: *DNMT3A*-N757K, *NPM1*-W288fs*12, *IDH1*-R132H, *NRAS*-G12C, *FLT3*-ITD. Dotted lines indicate thresholds for calling cells mutant or WT (Methods). Cells below the blue threshold are called 'WT'. Cells above the red threshold are called 'mutant'. Cells between the blue and the red threshold are called 'undetermined'. Only cells for which single-cell genotyping did not fail (Methods) are shown.

(B) As in (A), but from coding DNA (cDNA) amplicons for three loci in pt11 genotyped additionally from cDNA.

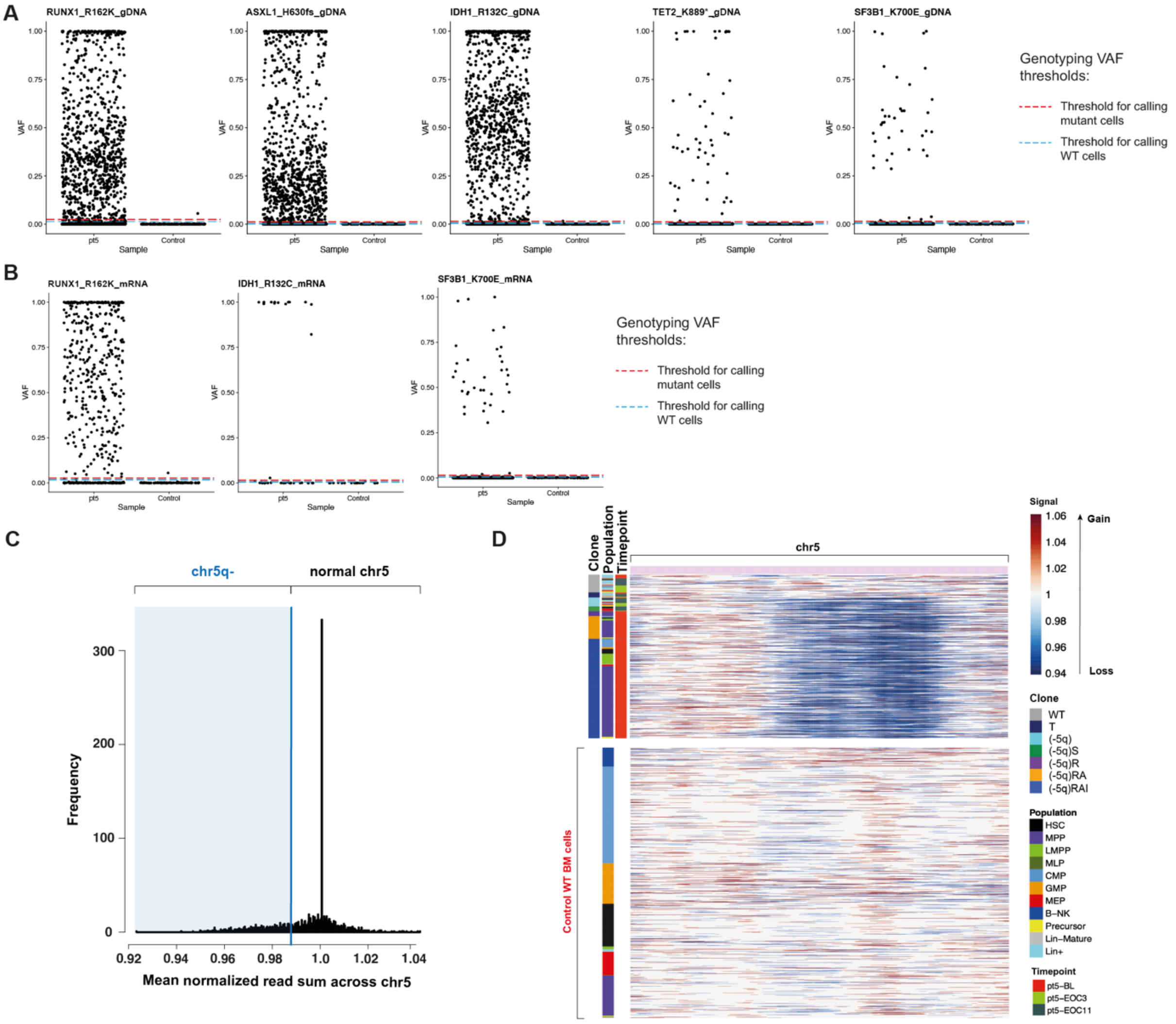


**Supplemental Figure 3. Single-cell genotyping and CNA calling for pt5.**

(A) As in Supplemental Figure 2A but from gDNA amplicons across five profiled mutant loci in pt5. The mutations are: *RUNX1*-R162K, *ASXL1*-H630fs, *IDH1*-R132C, *TET2*-K889*, *SF3B1*-K700E.

(B) As in (A), but from cDNA amplicons across the three mutant loci in pt5 genotyped additionally from cDNA.

(C) Histograms of mean normalized read sum across chromosome 5 (the first 30% of genes across chromosome 5 were excluded; detailed in Methods) for each cell (either from pt5 or from a healthy control BM). Cells displaying values below the blue threshold are called chr5q- (i.e. they have del(5)(q22q35), which was identified by karyotyping).

(D) Heatmaps showing RNA expression across chromosome 5 (x-axis) in single cells (y-axis) for pt5 (above; n = 728 cells) and a healthy control BM (below; n = 1197 cells). The bars to the left indicate the clone, the cell population, and the time point for each cell. Clone color-coding as in Figure 3A. On the heatmap, red indicates higher and blue indicates lower signal. Only cells passing scRNA-seq QC are shown.

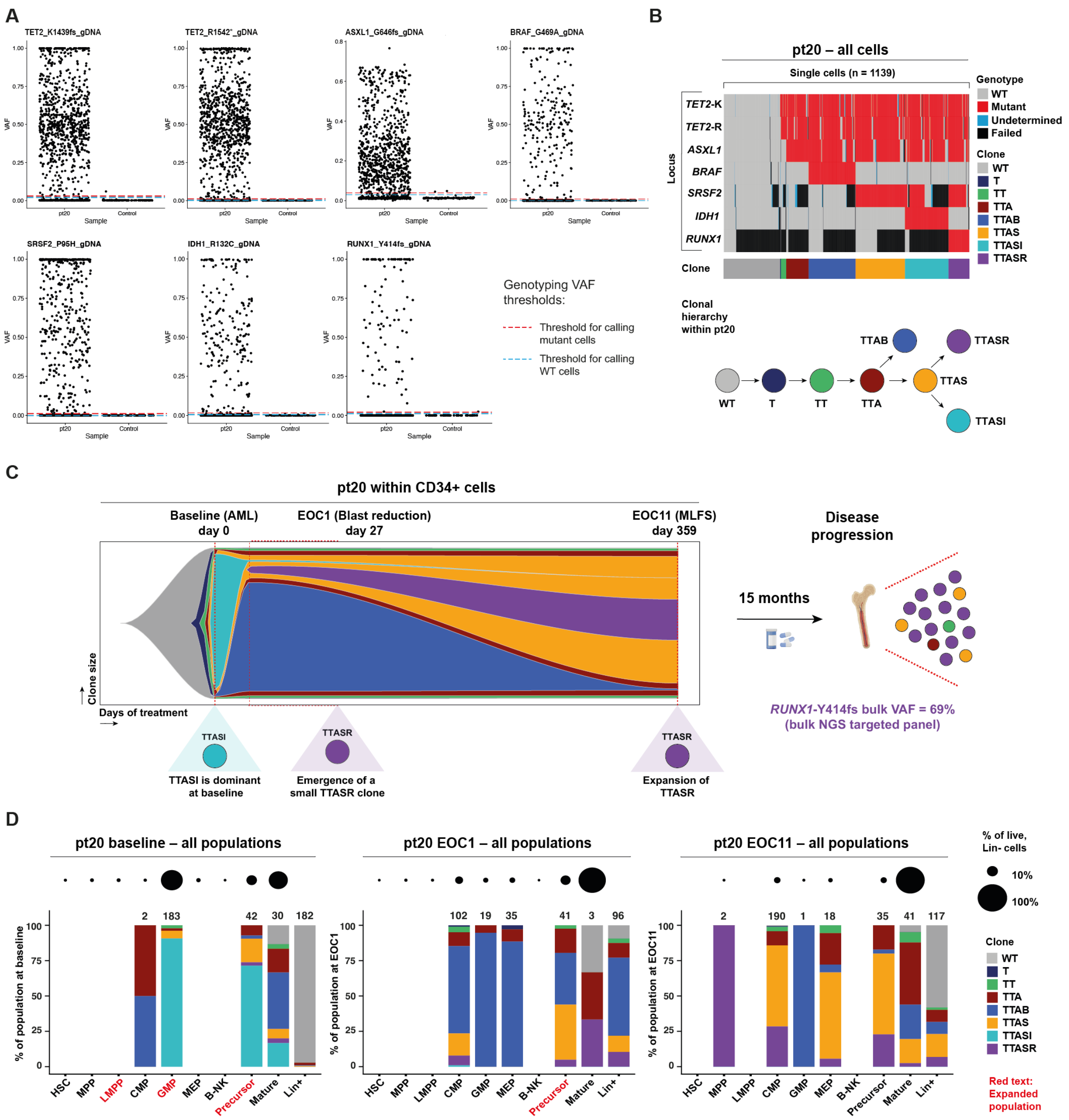


**Supplemental Figure 4. In pt20, a resistant *RUNX1*-mutated clone emerges 25 months prior to treatment failure.**

(A) As in Supplemental Figure 2A but from gDNA amplicons across seven profiled mutant loci in pt20. The mutations are: *TET2*-K1439fs*9, *TET2*-R1543*, *ASXL1*-G646fs*11, *BRAF*-G469A, *SRSF2*-P95H, *IDH1*-R132C, and *RUNX1*-Y414fs*186.

(B) As in Figure 2A but for pt20 (n = 1139). Genotyped mutations as in (A). *TET2*-K indicates *TET2*-K1439fs*9 and *TET2*-R indicates *TET2*-R1543*. Clone abbreviations are: WT: wild-type (gray). T: *TET2*-K mutant (dark blue). TT: *TET2*-K and *TET2*-R mutant (green). TTA: *TET2*-K, *TET2*-R, and *ASXL1* mutant (brown). TTAS: *TET2*-K, *TET2*-R, *ASXL1*, and *SRSF2* mutant (orange). TTASI: *TET2*-K, *TET2*-R, *ASXL1*, *SRSF2*, and *IDH1* mutant (cyan). TTAB: *TET2*-K,

*TET2*-R, *ASXL1*, and *BRAF* mutant (blue). TTASR: *TET2*-K, *TET2*-R, *ASXL1*, *SRSF2*, and *RUNX1* mutant (purple).

(C) As in Figure 2B but within $CD34^+$ HSPCs in pt20 (no enrichment for specific populations was performed), showing the baseline, EOC1, and EOC11 time points. Clone color-coding as in (B). Cell numbers are: baseline: 185; EOC1: 156; EOC11: 211. AML, *de novo* AML; MLFS, morphological leukemia-free state. Right, 15 months following EOC11, pt20 progressed and the VAF of the *RUNX1*-Y414fs*186 mutation was 69% (Supplemental Table 1).

(D) As in Figure 2C but for pt20 at baseline (left), EOC1 (middle), and EOC11 (right) time points. Clone color-coding as in (B).

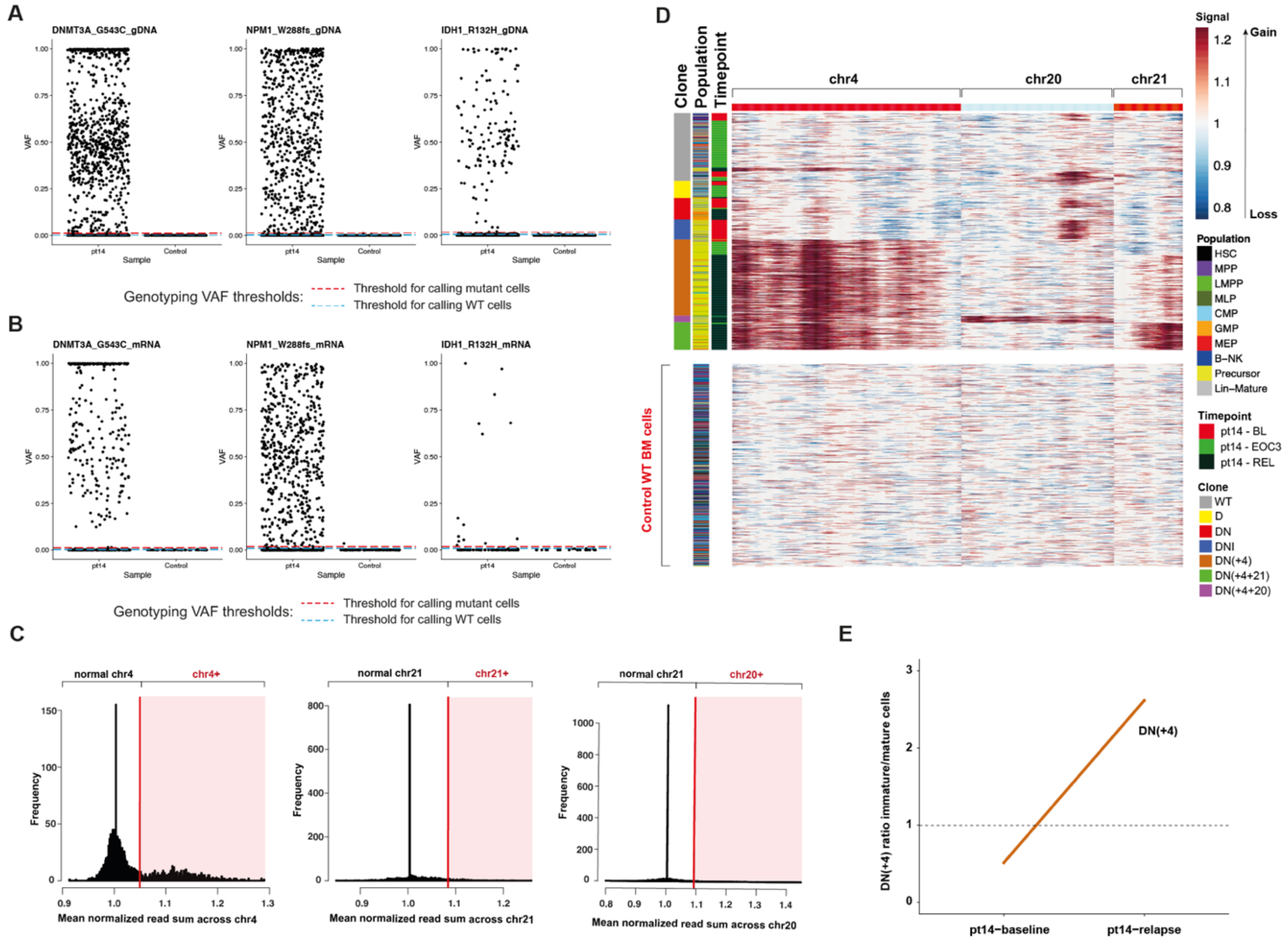


**Supplemental Figure 5. Single-cell genotyping and CNA calling for pt14.**

(A) As in Supplemental Figure 2A but from gDNA amplicons across three profiled mutant loci in pt14. The mutations are: *DNMT3A*-G543C, *NPM1*-W288fs*12, *IDH1*-R132H.

(B) As in (A), but from cDNA amplicons.

(C) Histograms of mean normalized read sum across chromosomes 4, 21, and 20 (genes across the entire chromosomes were used) for each cell (either from pt14 or from a healthy control BM). Cells displaying values above the red thresholds are called chr4+ (i.e. they have +4), chr21+ (i.e. they have +21), and chr20+ (i.e. they have +20). In pt14, +4, +20, and +21 were identified by karyotyping.

(D) Heatmaps as in Supplemental Figure 3D but for pt14 (above; n = 1251 cells) and a WT healthy control BM (below; n = 1057 cells) across chromosomes 4, 20, and 21 (x-axis; chromosomes indicated above) in single cells (y-axis). Clone color-coding as in Figure 4A.

(E) Line graph showing the ratio of the DN(+4) clone frequency in immature (HSPC and GM Precursor) versus mature myeloid cells, at baseline and relapse. A ratio above 1 (dashed line) indicates higher DN(+4) clonal frequency in immature cells; a ratio below 1 indicates higher DN(+4) clonal frequency in mature myeloid cells. The estimated clonal frequency in HSPC was corrected for FACS enrichment of HSPC sub-populations.

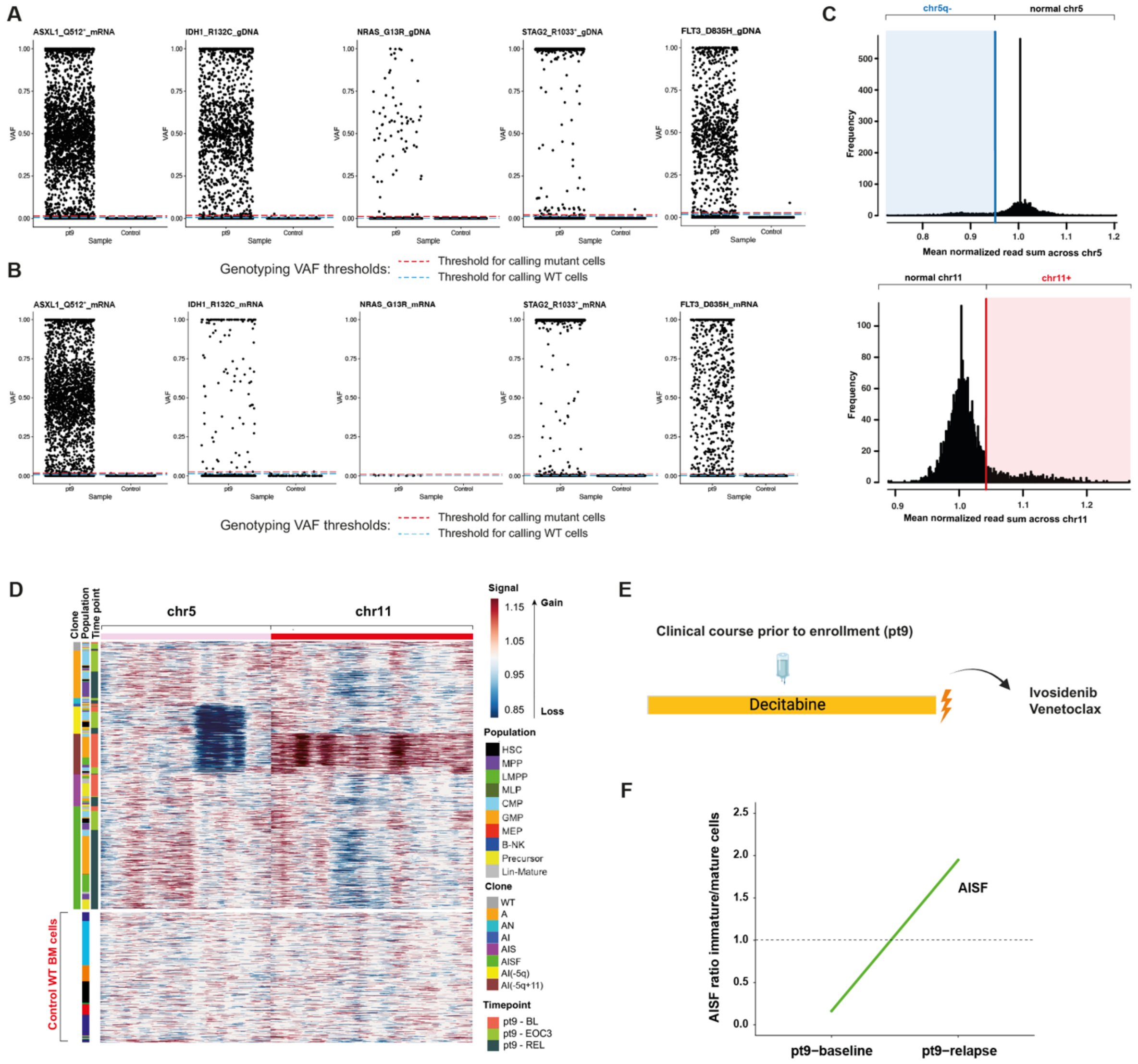


**Supplemental Figure 6. Single-cell genotyping and CNA calling for pt9.**

(A) As in Supplemental Figure 2A but from gDNA amplicons across five profiled mutant loci in pt9. The mutations are: *ASXL1*-Q512*, *NRAS*-G13R, *IDH1*-R132C, *STAG2*-R1033*, *FLT3*-D835H.

(B) As in (A), but for cDNA amplicons.

(C) Histograms of mean normalized read sum across chromosome 5 (left; the last 50% of genes across chromosome 5 were used; detailed in Methods) or chromosome 11 (right; the whole chromosome was used), for each single cell (either from pt9 or from a healthy control BM). Top, cells displaying values below the blue threshold are called chr5q- (i.e. they have del(5)(q22q33)). Bottom, cells displaying values above the red threshold are called chr11+ (detailed in Methods). In pt9, del(5)(q22q33) and +11 were identified by karyotyping.

(D) Heatmaps as in Supplemental Figure 3D but for pt9 (above; n = 2340 cells) and a healthy control BM (below; n = 1057 cells) across chromosomes 5 and 11.

(E) pt9 was enrolled after relapsing under decitabine treatment.

(F) As in Supplemental Figure 5E but for the AISF clone in pt9.

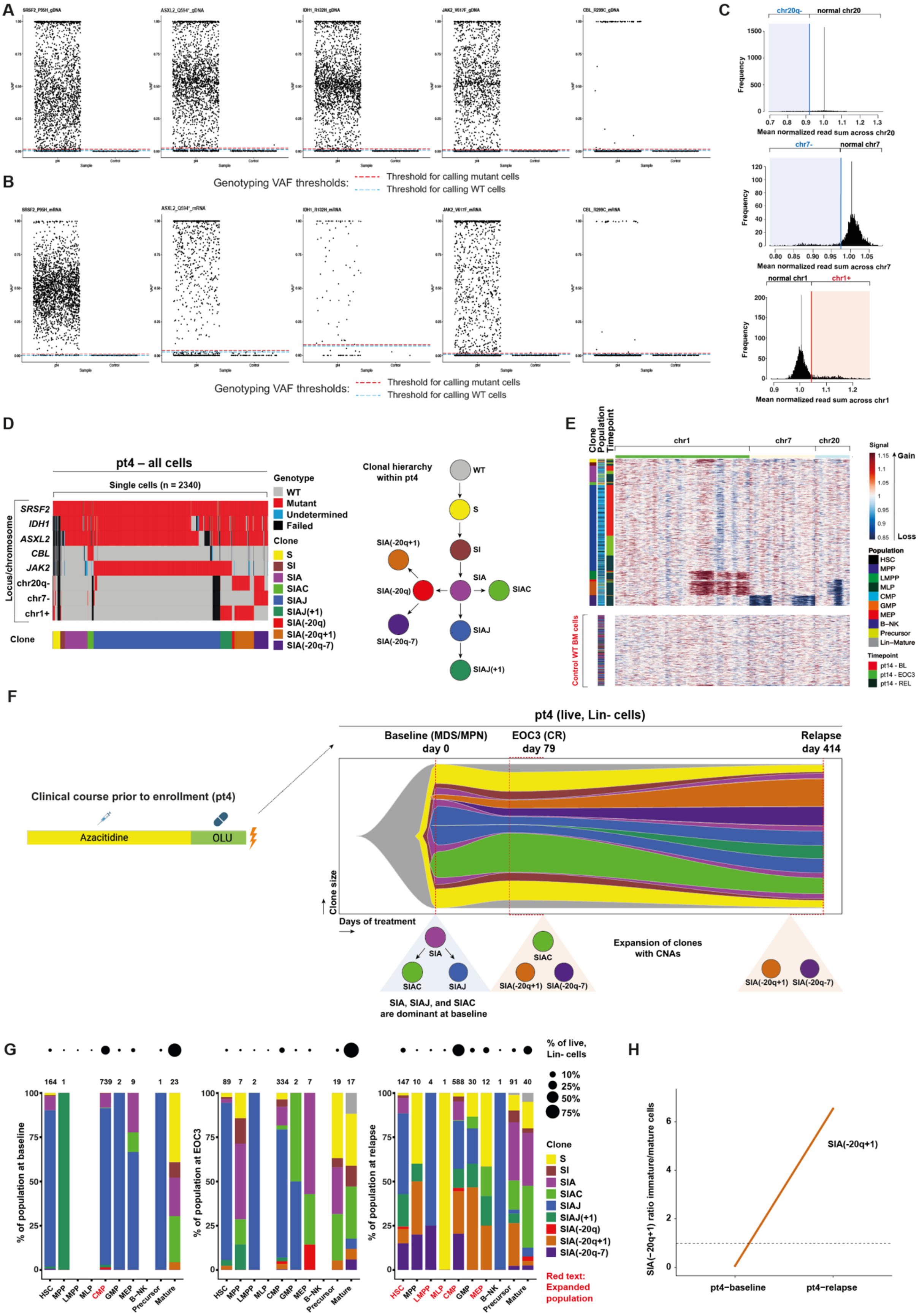
A
SRSF2_P95H_gDNA
ASXL2_Q594*_gDNA
IDH1_R132H_gDNA
JAK2_V617F_gDNA
CBL_R299C_gDNA
B
Genotyping VAF thresholds:
Threshold for calling mutant cells
Threshold for calling WT cells
SRSF2_P95H_mRNA
ASXL2_Q594*_mRNA
IDH1_R132H_mRNA
JAK2_V617F_mRNA
CBL_R299C_mRNA
Genotyping VAF thresholds:
Threshold for calling mutant cells
Threshold for calling WT cells
C
chr20q-
normal chr20
Frequency
Mean normalized read sum across chr20
chr7-
normal chr7
Mean normalized read sum across chr7
normal chr1
chr1+
Mean normalized read sum across chr1
D
pt4 – all cells
Single cells (n = 2340)
Locus/chromosome
SRSF2
IDH1
ASXL2
CBL
JAK2
chr20q-
chr7-
chr1+
Clone
Genotype
WT
Mutant
Undetermined
Failed
Clone
S
SI
SIA
SIAC
SIAJ
SIAJ(+1)
SIA(-20q)
SIA(-20q+1)
SIA(-20q-7)
Clonal hierarchy within pt4
WT
S
SI
SIA
SIAC
SIAJ
SIAJ(+1)
SIA(-20q)
SIA(-20q+1)
SIA(-20q-7)
E
Clone
Population
Timepoint
chr1
chr7
chr20
Control WT BM cells
Signal
Gain
Loss
Population
HSC
MPP
LMPP
MLP
CMP
GMP
MEP
B-NK
Precursor
Lin-Mature
Timepoint
pt14 - BL
pt14 - EOC3
pt14 - REL
F
Clinical course prior to enrollment (pt4)
Azacitidine
OLU
pt4 (live, Lin- cells)
Baseline (MDS/MPN)
day 0
EOC3 (CR)
day 79
Relapse
day 414
Clone size
Days of treatment
SIA
SIAC
SIAJ
SIA, SIAJ, and SIAC are dominant at baseline
SIAC
SIA(-20q+1)
SIA(-20q-7)
Expansion of clones with CNAs
SIA(-20q+1)
SIA(-20q-7)
G
% of population at baseline
164 1 739 2 9 1 23
% of population at EOC3
89 7 2 334 2 7 19 17
% of population at relapse
147 10 4 1 588 30 12 1 91 40
HSC
MPP
LMPP
MLP
CMP
GMP
MEP
B-NK
Precursor
Mature
% of live, Lin- cells
10%
25%
50%
75%
Clone
S
SI
SIA
SIAC
SIAJ
SIAJ(+1)
SIA(-20q)
SIA(-20q+1)
SIA(-20q-7)
Red text: Expanded population
H
SIA(−20q+1) ratio immature/mature cells
SIA(-20q+1)
pt4−baseline
pt4−relapse

**Supplemental Figure 7. In pt4, differentiation delay of 2 resistant clones harboring chromosomal aberrations occurs during treatment.**

(A) As in Supplemental Figure 2A but from gDNA amplicons across five profiled mutant loci in pt4. The mutations are: *SRSF2*-P95H, *ASXL2*-Q594*, *IDH1*-R132H, *JAK2*-V617F, *CBL*-R299C.

(B) As in (A), but from cDNA amplicons.

(C) Histograms of mean normalized read sum across chromosomes 20 (top), 7 (middle), and 1 (bottom; the last 50% of genes across chromosome 1 were used; detailed in Methods) for each single cell (either from pt4 or from a healthy control BM). For chromosomes 20 and 7, cells displaying values below the blue thresholds are called chr20q- (i.e. they have del(20)(q11.2q13.3); above) and chr7- (middle), respectively. For chromosome 1, cells displaying values above the red threshold are called chr1+ (i.e. they have +1; bottom). In pt4, +1 and del(20)(q11.2q13.3) were identified by karyotyping. Due to a strong CNA signal in inferCNV (see panel (E)), chromosome 7 was also studied.

(D) As in Figure 2A but for pt4 (n = 2340 cells). The genotyped mutations as in (A). Clone abbreviations are: WT: wild-type (gray). S: *SRSF2* mutant (yellow). SI: *SRSF2* and *IDH1* mutant (brown). SIA: *SRSF2*, *IDH1*, and *ASXL2* mutant (light purple). SIAC: *SRSF2*, *IDH1*, *ASXL2*, and *CBL* mutant (light green). SIAJ: *SRSF2*, *IDH1*, *ASXL2*, and *JAK2* mutant (blue). SIAJ(+1): *SRSF2*, *IDH1*, *ASXL2*, and *JAK2* mutant with chr1+ (dark green). SIA(-20q): *SRSF2*, *IDH1*, and *ASXL2* mutant with chr20q- (red). SIA(-20q+1): *SRSF2*, *IDH1*, and *ASXL2* mutant with chr20q- and chr1+ (brown). SIA(-20q-7): *SRSF2*, *IDH1*, and *ASXL2* mutant with chr20q- and chr7- (dark purple).

(E) Heatmaps as in Supplemental Figure 3D but for pt4 (above; n = 2233 cells) and a healthy control BM (below; n = 1057 cells) across chromosomes 1, 7, and 20 (x-axis; chromosomes indicated above) in single cells (y-axis). Clone color-coding as in (D).

(F) Left, pt4 was pre-treated first with azacitidine monotherapy and then with olutasidenib (OLU); subsequently, the patient was enrolled in the trial having failed both treatments. Right, as in Figure 2B but for pt4. Clone color-coding as in (D). Inferred clone sizes in Supplemental Table 4.

(G) As in Figure 2C but for pt4. Clone color-coding as in (D).

(H) As in Supplemental Figure 5E but for the SIA(-20q+1) clone in pt4.

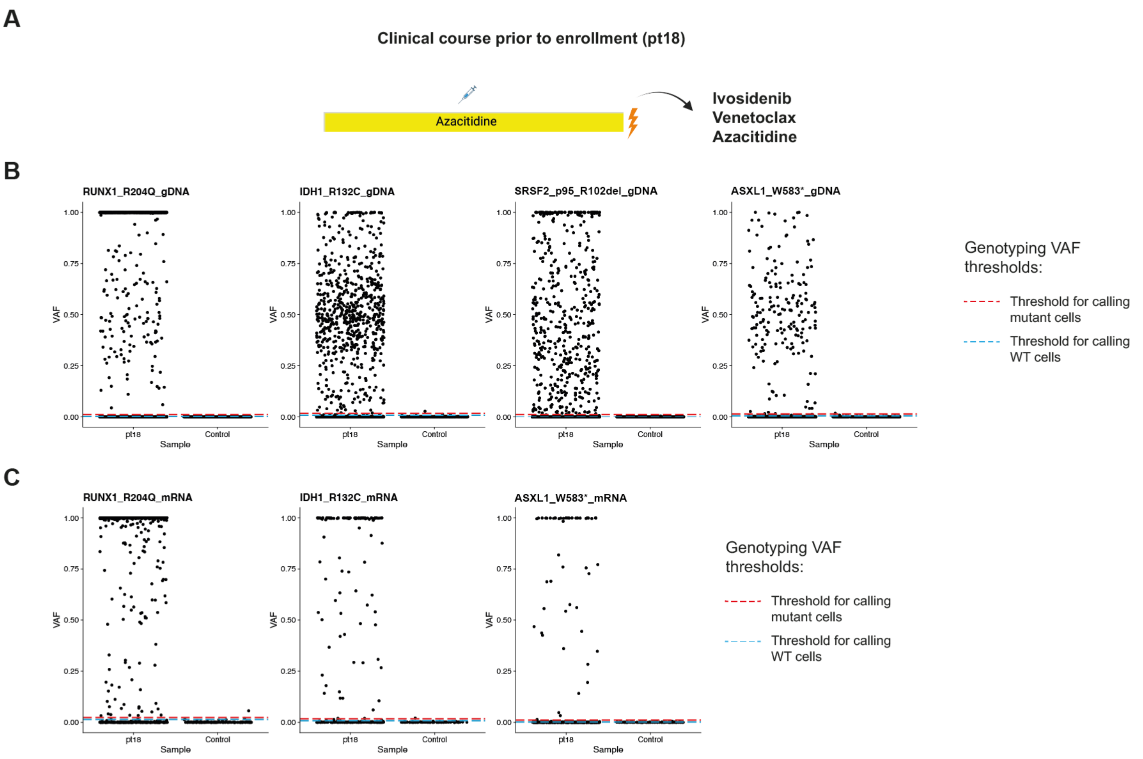


**Supplemental Figure 8. Single-cell genotyping for pt18.**

(A) pt18 relapsed after AZA monotherapy prior to enrollment to the trial.

(B) As in Supplemental Figure 2A but from gDNA amplicons across four profiled mutant loci in pt18. The mutations are: *RUNX1*-R204Q, *IDH1*-R132H, *SRSF2*-P95_R102del, *ASXL1*-W583*.

(C) As in (A), but from cDNA amplicons for three loci genotyped additionally from cDNA.

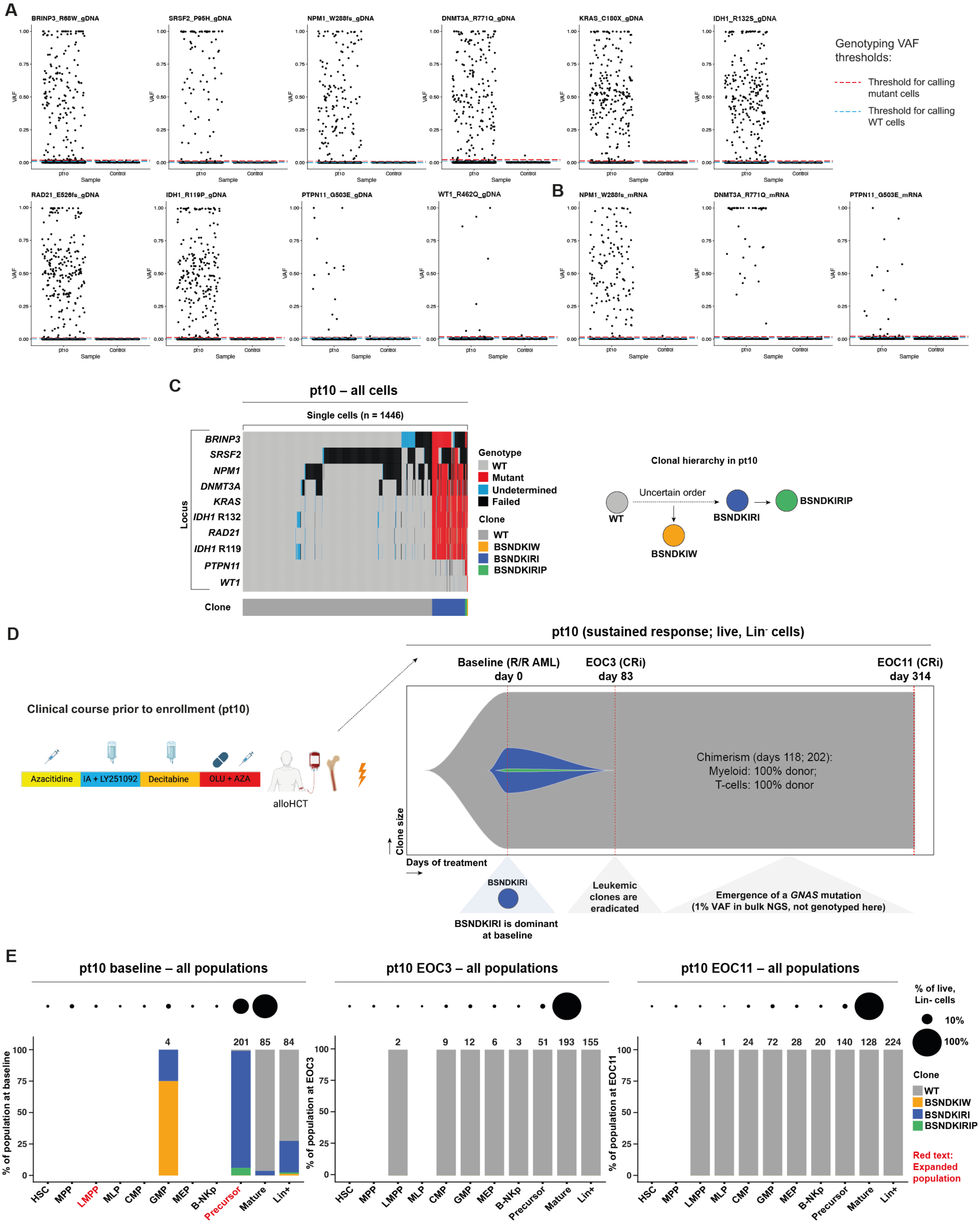


**Supplemental Figure 9. In pt10, sustained response is associated with clearance of all leukemic clones.**

(A) As in Supplemental Figure 2A but from gDNA amplicons across ten profiled mutant loci in pt10. The mutations are: *BRINP3*-R68W, *SRSF2*-P95H, *NPM1*-W288fs*12, *DNMT3A*-R771Q,

*KRAS*-C180X, *IDH1*-R132S, *RAD21*-E526fs*9, *IDH1*-R119P, *PTPN11*-G503E, and *WT1*-R462Q.

(B) As in (A), but from cDNA amplicons across three loci in pt10 genotyped additionally from cDNA.

(C) As in Figure 2A but for pt10 (n = 1446 cells). The genotyped mutations are as in (A). Clone abbreviations are: WT: wild-type (gray). BSNDKIW: *BRINP3*, *SRSF2*, *NPM1*, *DNMT3A*, *KRAS*, *IDH1* (R132S), and *WT1* mutant (orange). BSNDKIRI: *BRINP3*, *SRSF2*, *NPM1*, *DNMT3A*, *KRAS*, *IDH1* (R132S), *RAD21*, and *IDH1* (R119P) mutant (blue). BSNDKIRIP: *BRINP3*, *SRSF2*, *NPM1*, *DNMT3A*, *KRAS*, *IDH1* (R132S), *RAD21*, *IDH1* (R119P), and *PTPN11* mutant (green). The order of mutation acquisition prior to the BSNDKIRI clone was unclear.

(D) Left, pt10 was pre-treated with 4 lines of therapy: (a) azacitidine (AZA); (b) idarubicin, cytarabine (IA), and the CXCR4 peptide antagonist LY2510924; (c) decitabine; and (d) olutasidenib (OLU) with AZA. Subsequently, pt10 underwent allo-HSCT. Following relapse post-allo-HSCT, the patient was enrolled in the trial. Right, as in Figure 2B but for pt10, showing the baseline, EOC1, and EOC11 time points. Clone color-coding as in (C). A *GNAS*-R201H mutation, not detected at baseline, was detected at remission at 1% VAF via the bulk targeted NGS panel but was not genotyped by TS+. Inferred clone sizes in Supplemental Table 4.

(E) As in Figure 2C but for pt10, at baseline (left), EOC1 (middle), and EOC11 (right) time points. Clone color-coding as in (C).

**Supplemental Tables Titles**

**Supplemental Table 1. Patient Clinical Profiles, NGS Targeted Panels, and Genetic and Cytogenetic Changes at Times of Sampling**

**Supplemental Table 2. Flow Cytometry Analysis of Normal BM Donors (from the MARCH Cohort; Jakobsen et al., 2024) and Patient Samples.**

**Supplemental Table 3. TARGET-seq+ Primers and Library Success Rates**

**Supplemental Table 4. Estimated Clone Sizes and Confidence Intervals in All Live, Lin$^-$ Cells**
